# Multiplexed drug testing of tumor slices using a microfluidic platform

**DOI:** 10.1101/2020.02.25.965137

**Authors:** L. F. Horowitz, A.D. Rodriguez, Z. Dereli-Korkut, R. Lin, K. Castro, A. Mikheev, R.J. Monnat, A. Folch, R.C. Rostomily

## Abstract

Current methods to assess the drug response of individual human cancers are often inaccurate, costly, or slow. Functional approaches that rapidly and directly assess the response of patient cancer tissue to drugs or small molecules offer a promising way to improve drug testing, and have the potential to identify the best therapy for individual patients. We developed a digitally-manufactured microfluidic platform for multiplexed drug testing of intact cancer slice cultures, and demonstrate the use of this platform to evaluate drug responses in slice cultures from human glioma xenografts and patient tumor biopsies. This approach retains much of the tissue microenvironment and can provide results rapidly enough, within days of surgery, to guide the choice of effective initial therapies. Our results establish a useful preclinical platform for cancer drug testing and development with the potential to improve cancer personalized medicine.

## INTRODUCTION

Despite advances in targeted and immune therapies, cancer treatment continues to face enormous challenges as it moves toward the goal of rationally chosen, personalized therapy ^1, 2^. Critically, while molecular features alone can guide treatment, they cannot reliably predict an individual patient’s functional response to targeted, immune, or conventional treatment. However, *in vitro* functional tests on an individual’s cancer could help to predict that patient’s outcome, even without any molecular knowledge. In addition, the high cost of drug development is slowing the pace of discovery. Despite success in identifying drugs or small molecules that target specific mutations and pathways^3^ the majority of cancer patients do not currently qualify for genomically targeted drug treatment and responses are limited by cell intrinsic resistance and microenvironmental factors^2^. Furthermore, it has become clear that the presence of a targetable or ‘actionable’ mutation does not guarantee the success of drug treatment^2^, and may lead to substantially different outcomes for different patients and for different cancers (e.g. the effective targeting of the BRAF V600E mutation in melanoma but not in colon cancer)^4^.

Functional approaches that directly assess the response of patient cancer tissue to drugs or drug combinations offer a promising way to improve drug testing and have the potential to rapidly identify the best therapy for individual patients^2^. Functional assays can potentially complement and extend genomics-based approaches by capturing key determinants of therapeutic response such as tissue architecture, tumor heterogeneity, and the tumor microenvironment^2^. Hence these assays promise to improve both the speed and success of drug development by offering a more physiologically-relevant human drug testing platform to complement animal testing; presently each drug may take up to a decade to move to clinical application and at a substantial cost (estimated at ∼650 million dollars/drug)^5^, in large part due to high failure rates.

Diverse functional assay platforms have been developed that assess drug responses in tumor tissue samples. Early strategies that used dissociated 2D cell cultures (called “Cell Culture Drug Resistance Testing” or CCDRT) from patient tumors were unreliable at predicting therapy outcomes, and have been abandoned in favor of 3D tissue models^6^. The shortcomings of the 2D formats also highlighted the emerging recognition of the importance of the tumor microenvironment in regulating malignancy and chemosensitivity ^7^. The 3D platforms attempt to capture both the architecture and microenvironment in order to more closely resemble the primary tumor. These approaches include organoid cultures, micro-dissected tumor spheroids, tumor slices, patient-derived xenograft (PDX) mouse models, and direct microdelivery to patient tumors^2^. While microneedles or microdevices that deliver drugs into patient tumors *in vivo* offer maximal preservation of the tumor microenvironment, issues of tumor accessibility and patient safety are likely to limit their application^8, 9^. PDX models permit the study of drug responses in an intact organism, including immune checkpoint blockade in humanized PDX^10^, but the rest of the microenvironment is from the host mouse. While syngeneic and transgenic mouse tumor models offer a native and intact microenvironment – key for studies of immune-related therapies – both the tumor and the microenvironment are completely non-human^11^. Importantly, PDX from individual patients cannot be grown rapidly enough to inform initial post-operative therapeutic decisions.

Like tumor spheroids^12, 13^, but to a greater extent, tumor-derived slice cultures retain and sample the original tumor’s content and 3D structure, including ECM, non-tumor stromal cell types, and biochemical pathways^14–16^. The implication of various cellular (e.g. fibroblasts, lymphocytes, macrophages, and endothelial cells), matrix, and metabolic components of the tumor microenvironment as drivers of drug responsiveness highlights the importance of retaining these elements in a pre-clinical screening platform ^17, 18^. Drug testing has been performed on slice cultures from multiple tumor types, including brain, breast, GI, skin, and pancreas tumors ^14, 15, 19–21^. Slice cultures have the potential to improve functional drug screens by allowing the rapid testing of different potential therapies. For example, drug testing of slice cultures from PDX or from primary tumors has been shown to correlate with treated responses in patients with pancreatic^21^ and breast cancers^22^. However, in most studies with slice culture, drug is uniformly applied to the slice. As the number of slices per tumor is limited, the numbers of conditions that can be tested are limited.

The combination of microfluidic drug delivery with slice culture has the potential to enable multiplexed drug delivery and response assessment while preserving the native microenvironment and tissue microarchitecture. PDMS-based microfluidic perfusion devices for cultured tissue slices have been developed that perform electrophysiological recordings to brain slices^23, 24^, multiplexed drug delivery to normal brain slices^25^, and drug delivery to lymph node slices^26^. Finally, a 5-channel microfluidic device was used to trap and culture mini-discs punched out from tumor slices (one inside each closed channel), and for one patient’s sample, to test responses to a single drug^27^.

We present here an intuitive, multi-well based microfluidic platform to test responses to multiple drugs on individual cancer slice cultures. The microfluidic device is digitally manufactured by lasercutting in poly(methyl methacrylate) (PMMA). This PMMA device builds upon our prior work with a PDMS-based microfluidic device^25^. Both share a higher throughput, user-friendly format: an accessible open culture surface, multiple underlying fluidic channels (40 here, 80 in the prior PDMS device, and only up to 10 channels for the other devices above), easy solution delivery from open wells, and a single output channel. However, this PMMA device offers two critical improvements. First, the faster and cheaper manufacture of the PMMA device by digital design and laser cutting requires much less specialized labor than does the complex manufacture of the earlier multi-layer PDMS device by photolithography. Second, because drug absorption is a concern with PDMS ^28^, but not with PMMA (a solid plastic), PMMA may be more appropriate for drug studies, although both materials can have drug adsorption on the surface^29, 30^ This 40 channel PMMA device allows for the multiplexed testing of at least 20 drug conditions. We demonstrate the use of this device for the multiplexed delivery of drugs and dyes to xenograft-derived human glioblastoma (GBM) slices in culture. We also demonstrate differences in drug responses for GBM cells grown in 2D culture versus in tumor slice culture, as well as for xenograft tumor slices derived from flank versus intracranial tumors. These differences emphasize the importance of tumor structure and microenvironment in order to accurately assess therapeutic response. Finally, we demonstrate the use and versatility of this microfluidic platform by assessing drug responses in primary, patient-derived tumor slices from GBM and colorectal cancer (CRC) liver metastases.

## RESULTS

### Development and characterization of a microfluidic device for drug delivery to tumor tissues

Fig. 1a outlines our approach to evaluating drug responses of tumor-derived slice cultures, both on and off our microfluidic device. We used xenograft tumors for the initial studies and patient samples for more advanced studies. Xenograft tumors provide the biological reproducibility needed for the initial studies. We generated flank or intracranial xenograft tumors by injection of GBM cell lines (U87 or GBM8) into immunodeficient nude mice (*Foxn1^nu^*). Tumor slices were cut using a vibratome (∼250 µm thick) and grown as organotypic slice cultures on a polytetrafluoroethylene (PTFE) porous membrane with an air interface above and medium below. Drugs were applied to the bottom surface in the medium, either as single drugs per well for off-device well experiments or as stripes of multiple drugs for the microfluidic device. For live/dead analysis on intact slices, we imaged the bottom surface for fluorescent nuclear dye markers. For histology and immunostaining (cell death, cell types), we analyzed thin cross-sections of the slices. We chose standard imaging (epifluorescence over confocal) and histology approaches that maximize the ease and accessibility of our analyses.

The operation of the microfluidic device was designed to be intuitive to untrained users. Two laser-cut PMMA layers create a device with a multi-well plate at the top and a microchannel network layer below (Fig. 1b and c, Supplemental Fig. S1). PMMA plastic is optically clear and biocompatible^31^. The multi-well plate containing 40 input wells (1.6 mL each) is bonded to the microchannel network that connects each well to the central culture area of the device. The overall dimensions and number of turns are the same for all channel paths to ensure similar resistance, and thus similar flow rates for all channels, during flow. A binary arbor combines all 40 channels into a single output. Flow is established on all channels simultaneously by applying suction at the outlet at a low flow rate by means of a syringe pump, which draws from all 40 wells at approximately the same rate. The user simply places the tissue slice and membrane into the central culture area of the device, fills the wells with a pipet as with any multi-well plate, and runs the device with a single syringe pump. In the center portion of the device (2 cm × 1 cm), the microchannels (∼140 µm-wide at the top) have open roofs and are spaced out 500 µm from center to center. When the tissue slice on a porous culture membrane is placed on top of the open-roof microchannels, the membrane becomes the roof of these channels and permits fluid flow underneath the slices within the channels when suction is applied, the membrane must completely cover the open channel windows in order to create a proper seal between the membrane and the smooth surface surrounding each opening of the channels. Otherwise, unwanted suction of air would perturb the balance of flow in the other channels. The resistance of the input channels is high enough to prevent appreciable leakage of fluid for at least 15 minutes after the stoppage of negative flow. When solutions containing drugs flow under the membrane, the drugs move through the membrane and penetrate up into the tissue by diffusion. Tissues simply grow on top of the membrane as in culture, and a laser-cut lid covers the top to reduce evaporation and prevent the tissue from drying. While some slices form an attachment to the membrane in culture, tissue attachment is not necessary for running the device during which time negative pressure also holds the tissue/membrane in place. Alternating drug and buffer lanes act as a source and sink, respectively, to limit lateral diffusion along the membrane during drug application.

**Fig. 1:**
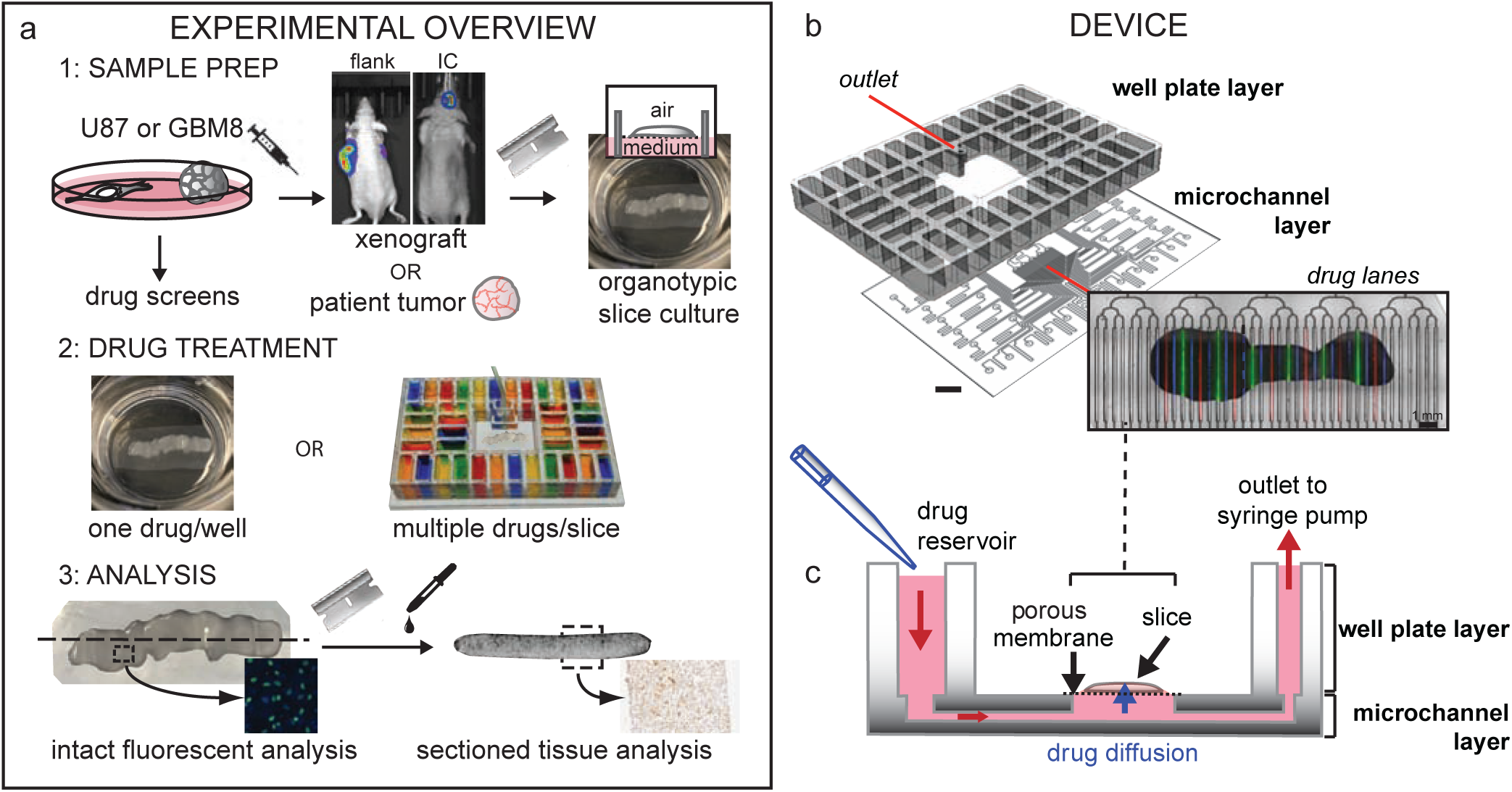
Oncoslice device for multiplexed microfluidic drug delivery to tissue. **a**) Slices from glioma cell line-derived xenografts or patient tumors cultured on porous membranes were treated with drugs in 6-well plates (1 drug/well) or on the microfluidic device. A cell line drug screen guided drug choice. Responses were measured by fluorescence or tissue staining. **b**) Oncoslice device assembly by bonding of laser-cut PMMA 40-well plate and microchannel layers. Image shows delivery of Cell Tracker Red, Cell Tracker Green, and Hoechst (blue) dyes to U87 xenograft slices. **c**) Device cross-section shows wells, microchannel delivery to tissue slices, and a single syringe pump outlet. Scale bar: 1 cm.

To visualize and quantify drug delivery into live tissue using the device, we analyzed cross-sections after drug treatment, and thus visualized the extent of drug delivery both laterally between lanes, and vertically from the drug source at the membrane side up to the surface. We were able to look at drug delivery directly by fluorescence after quick freezing tissue to prevent further diffusion, or indirectly by cell drug readouts such as immunostaining. We exposed U-87 MG (U87) glioma xenograft flank tumor tissue slices to two fluorescent chemicals with a molecular weight (M.W.) similar to that of most small molecule drugs (Fig. 2). Doxorubicin (DOX), an anthracycline chemotherapy drug used to treat a wide range of cancers, fluoresces red (M.W. 544 g/mol). Although DOX fluorescence can change with binding to biomolecules and with high concentrations over 45 µM^32^, we measure emission at a relatively linear range of concentrations (applied at 10 µM), and assume similar distribution of tissue biomolecules throughout the relatively homogeneous xenograft tumor tissue. Our second chemical, the nuclear dye Hoechst 33342, fluoresces blue upon binding DNA (M.W. 453 g/mol). We used fresh frozen cryosections that are cut orthogonal to the membrane plane (in order to expose the interior of the slice) after the slices are exposed to fluorescent drugs or chemicals. In our orthogonal cryosections, we can measure (without fluorophore spectrum overlap) the fluorescence from the two chemicals within the tissue using standard red and blue channels. When the two chemicals were applied in alternating lanes (without buffer lanes) for a short timepoint (4 h), both DOX and Hoechst distributed vertically into the tissue in a gradient as expected (Fig. 2a,c; over a 55 µm wide profile). Next, we measured the vertical diffusional spread of the chemicals within the tissue after running the device for 48 h, reflecting the longer periods necessary for drug testing (Fig. 2b,c; 55 µm wide profile). In this case, we spaced the repeats of a chemical 8 lanes apart (4 lanes between different chemicals). An overall increase in fluorescence over background throughout the depth of the tissue denotes an increase in vertical diffusional spread as compared to over 4 h. The increase in fluorescence seen on the top surface is likely due to concentration from evaporation that occurs at the air interface. Note that we do not directly convert fluorescence to concentration given the complexities of DOX and Hoechst fluorescence signals. While the M.W. of these two drugs is similar to that of other small-molecule drugs such as those used later in this study, and size (related to M.W.) is a primary determinant of diffusion, different molecules will have different biophysical properties, such as lipid solubility and binding, that may affect their distribution.

**Fig. 2:**
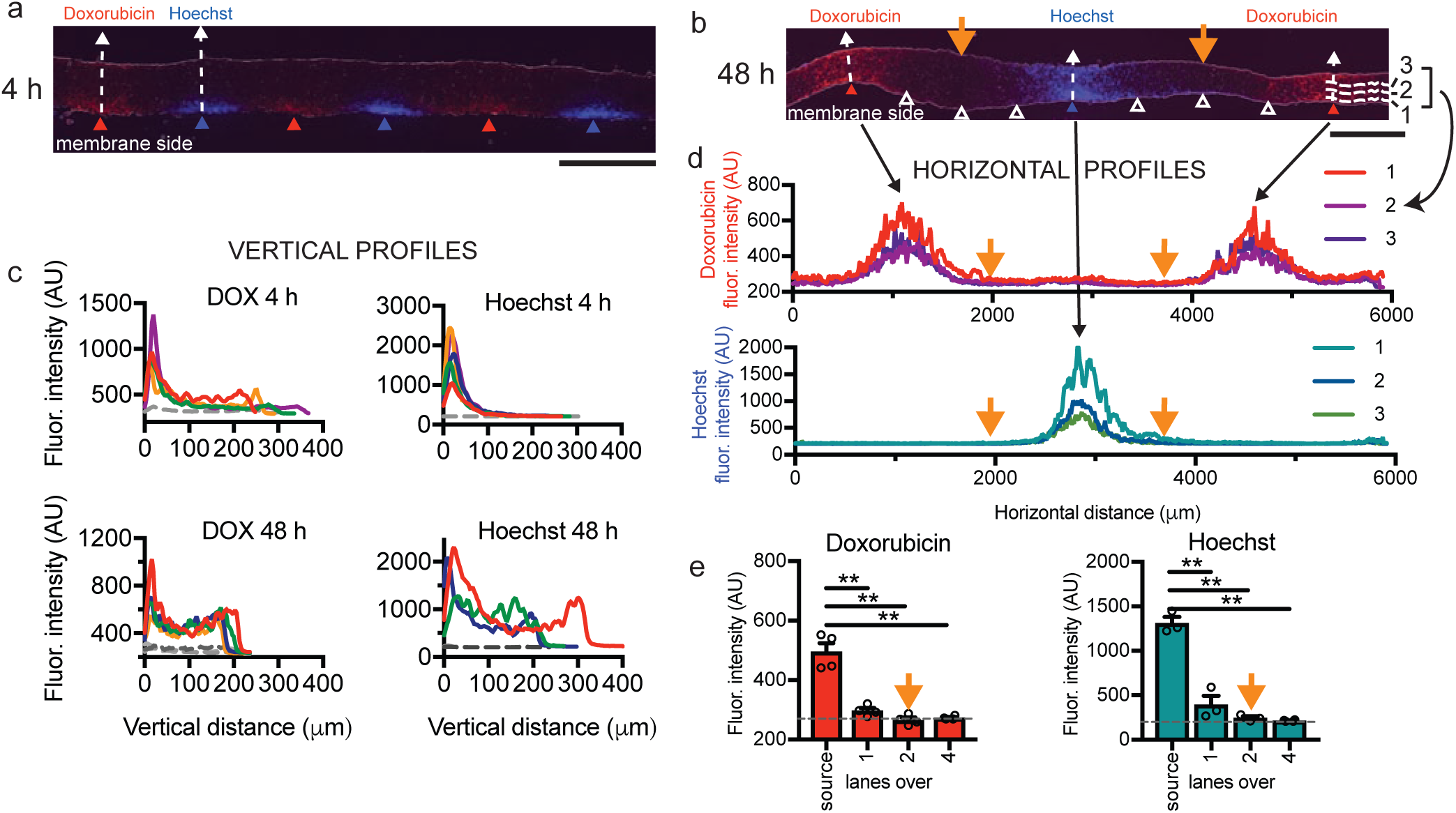
Fluorescent analysis for microfluidic drug delivery to tissue. Vertical concentration gradients of fluorescent doxorubicin (DOX, red) and Hoechst dye (blue) in U87 xenograft slices at 4 h (**a**) or 48 h (**b**), separated by buffer lanes (white triangles). **c**) Vertical fluorescence profiles above source lanes versus distance from the membrane at 4 h or 48 h. Colored lines represent average fluorescence at different lanes for 4 h (4 DOX, 5 Hoechst), and at spaced locations on 2 lanes for 48 h (4 DOX, 3 Hoechst). Gray profiles represent Hoechst for doxorubicin (and *vice versa*), averaged at 4 h and individual profiles at 48 h. **d**) Horizontal profiles at 48 h show lateral diffusion. Locations indicated in (**b**) (lines 1-3). **e**) Lateral fluorescence spread from source lanes at 48 h as indicated in (**d**). Average ± SEM, n=4 for DOX, n=3 for Hoechst except for 4 lanes over. One-way ANOVA, Tukey’s multiple comparison test. **p<0.0001. Scale bars: 500 µm (**a,b**).

We then assessed the conditions that would avoid cross-contamination between lanes. In order to determine whether our microfluidic device can deliver drugs applied two lanes apart without significant cross-contamination in the tissue directly above the delivery lanes, we examined the lateral diffusion of the fluorescent chemicals in the 48 h exposure experiment described above (Fig. 2b). DOX and Hoechst fluorescence signal clearly decreased with distance from the source lane, as quantitated with horizontal profiles (Fig. 2d). Close to the bottom surface (13-39 µm), we measured the signal over the estimated center of each lane (identified as the center of fluorescence), and then at ∼500 µm intervals of 52 µm width, Fig. 2e). As expected for one lane over from the dye source, we observed lower but significant signal (10%±4% for DOX and 16%±9% for Hoechst, ave±sem, n=3-4), measured as a percent of the source signal over the background signal (four lanes over). However, two lanes over where the next drug would be applied, we could not distinguish the signal from the background (−3%±4% for DOX and 0%±1% for Hoechst). Naturally, tissue consistency, experiment duration, and drug type all may affect the diffusion profiles, necessitating re-assessment of delivery for each tissue type and experimental setup. These experiments suggest that when we run our device with alternating source and buffer lanes, the regions above the source lanes remain relatively free from cross-contamination, and the regions above intervening buffer lanes would have contributions from their neighboring lanes. Thus for analysis of independent drug delivery, we may assess tissue directly above drug and vehicle control “source” lanes. Tissue above the intervening buffer “sink” lanes and the interchannel regions represent exposure to lower concentrations of drug.

### Temporal characterization of xenograft slice cultures and selection of drugs

Changes in cell health or composition that accompany adaptation of acutely isolated tumor slices to culture conditions over time could impact optimal timing and interpretation of dug screens. Therefore, prior to drug testing on slices, we performed baseline studies in which we characterized the slices cultured off-device. We first examined viability, growth and cell proliferation/death (Fig. 3a, Supplemental Fig. S2). Slices from U87 flank xenografts provided large, reproducible tumors well suited for device validation. First, we established that healthy slices could be maintained for a week with continued growth (Supplemental Fig. S2a-e). Proliferation (Ki-67 immunostaining), apoptosis (cleaved caspase-3, CC3, immunostaining), and non-specific cell death (Sytox Green, SG, dead nuclear stain) decreased over the first 1-3 days in culture (Fig. 3a, Supplemental Fig. S2f-k). As the tumor microenvironment may affect clinical responses to drugs, we documented the stromal cell population in the tumor slices by immunostaining (Fig. 3b, Supplemental Fig. S3). In flank tumors from U87-GFP xenografts, we quantitated the GFP+ U87 human tumor cells and non-human stromal cells derived from the mouse host using both GFP and immunostaining for human-specific HuNu antigen (Fig. 3a, Supplemental Fig. S3a-c). We estimated the total stromal cell population at between 10-20% of cells (Supplemental Fig. S3c). The presence of vascular endothelial cells (CD31+), immune cells (CD45+), macrophages (IBA-1+), and vimentin+ mesenchymal cells in the tumor stroma was confirmed by immunostaining (Fig. 3a, Supplemental Fig. S3a,b,d,e). In general, as quantified for IBA-1+ cells (Supplemental Fig. S3f,g), stromal cells gradually declined over the first 4 days in culture, except for a persistent, low level of vimentin+ cells of ∼0.2% between days 2-5. These cells are likely to be cancer-associated fibroblasts, but may represent endothelial cells or lymphocytes^33^ with atypical shapes. Therefore, for subsequent drug treatments, we chose a 48-hour window from day 1-3. At this time, slices have recovered from culture shock, and proliferation and stromal cells, thought to be therapeutically relevant features, are present.

**Fig. 3:**
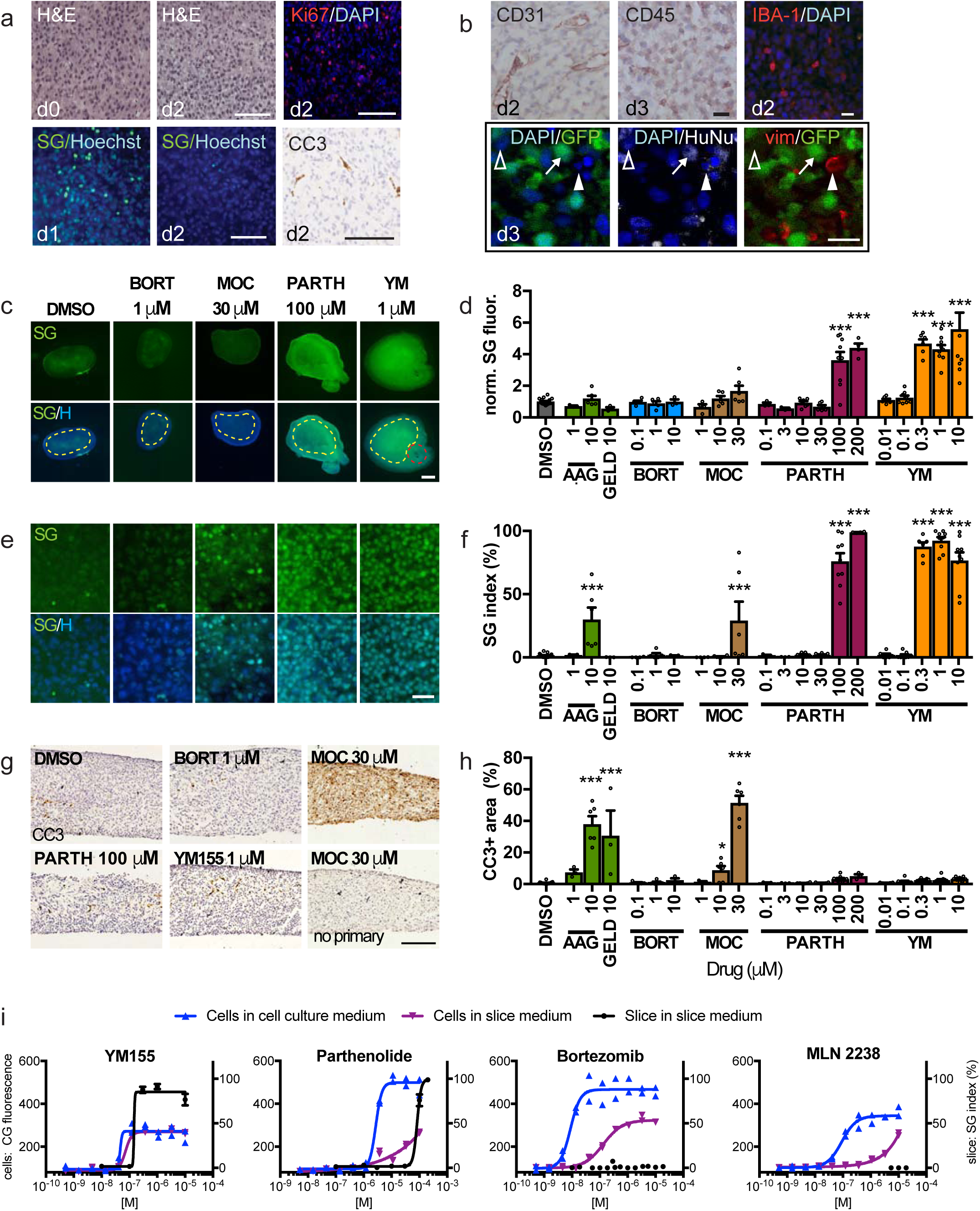
U87 glioma xenograft slice culture characterization and drug responses. **a)** Retention of tissue integrity in U87 flank xenograft slice cultures by H&E staining, cell proliferation detected by Ki-67 immunostaining (red, DAPI counterstain) and cell death by SYTOX Green (SG)/Hoechst (blue) staining or by cleaved-caspase 3 (CC3)/hematoxylin staining. **b)** Xenograft slices contain endothelial (CD31), immune (CD45) and macrophage (IBA-1) cells. Confocal images identify U87 glioma cells (arrow, GFP^+^/HuNu^+^), vimentin-positive mesenchymal stromal cells (vim, red, solid arrowhead) and additional non-tumor cells (open arrowhead). **c-f)** Slices treated for 2 or 3d and assessed for cell death by SG/Hoechst staining. Tanespimycin (AAG), geldanamycin, bortezomide (BORT, 3 d), mocetinostat (MOC, 3 d), parthenolide, and YM155 (YM). Cell death was quantified by mean fluorescence (dotted regions)(**c,d**) or by % SG^+^/total nuclei (**e,f**). Red outline in (**c**) is a positive control crush lesion. **g,h)** Apoptotic cell death quantified as CC3-stained area. No primary antibody control is shown. **i)** Cell death in U87 cells by loss of CellTox Green fluorescence (CG) when grown in conventional (blue) or slice (purple) medium, versus U87 xenograft slices (black line) in slice medium (SG index). For (**d,f,h**), average ± SEM; n= 17,3,6,5,5,3,3,4,6,6,3,3,7,6,9,4,6,8,6,9,9 slices for each drug condition, in order. One-way ANOVA versus DMSO with Dunnett’s multiple comparison test. *p<0.05, **p<0.001. Cell line assays were done in duplicate. Scale bars: 100 µm (**a,g**), 20 µm (**b,e**), 1 mm (**c**).

To select candidate drugs for testing on slices, we performed high-throughput drug screens in 2D cell culture with the GBM U87 and GBM8 glioma stem cell lines used to generate the xenograft tumors in this study. After a primary screen of a 350 anti-cancer drug library (Selleck Anti-Cancer Drug) (Supplemental Fig. S4a), we selected 32 drugs for secondary screens based on their clinical relevance, mechanisms of action, and activity in the two cell lines (Tables S1,2, Supplemental Fig. S4). For the screens, we assessed viability with the bioluminescent metabolic indicator CellTiter-Glo. For the secondary screen, we also assessed cell death with the fluorescent cell death marker, CellTox Green, because we performed slice culture response readouts with a cell death marker, Sytox Green, analogous to CellTox Green, instead of with a viability indicator. An example comparison of dose-responses to an individual drug (YM155), as well as overall determinations of drug activity using CellTiter-Glo and CellTox Green readouts, are shown in fig. S3B-E. The final drug set included cisplatin (CP; DNA/RNA synthesis inhibitor), tanespimycin (AAG; heat shock protein inhibitor), geldanamycin (GELD; autophagy/heat shock inhibitor), bortezomib (BORT; proteasome inhibitor), parthenolide (PARTH; NFkB inhibitor), YM155 (YM; E3 ligase/survivin inhibitor), mocetinostat (MOC; HDAC inhibitor) and MLN 2238 (MLN; a proteasome inhibitor).

### Cell death readouts for drug responses in U87 glioma xenograft slice cultures

The quantification and comparison of drug responses in slice cultures required development and optimization of quantitative drug response readouts. We used Sytox Green (SG), a fluorescent nuclear stain to detect non-specific cell death in intact tissue and cleaved caspase-3 (CC3) immunostaining to detect apoptotic cell death in tissue sections. One SG readout measured mean tissue fluorescence (normalized to vehicle control after subtraction of background) in low power images. The other SG readout, the SG index, measured SG+ dead green nuclei as a percentage of the total cell number (detected by the blue nuclear dye Hoechst) in high power images. For apoptosis, we measured the CC3+ area (as a percent of total tissue area). Note that these cell death measures do not differentiate between tumor cells and stromal cells. However, as we found above that slices contained approximately 90% tumor cells, any high percentage of cell death would most likely reflect mainly the death of tumor cells, with a potential small contribution from the stromal cells. We compared the results for each cell death readout using responses to 2 days of cisplatin administration in U87 flank xenograft slices off-device (Supplemental Fig. S5). All three methods yielded similar results, with significant increases in cell death between 30 and 100 µM (Supplemental Fig. S5).

Using these SG and CC3 readouts, we then compared dose responses to different drugs selected from the secondary screen in U87 flank-derived xenograft slice cultures off-device (Fig. 3c-h). While the two SG readouts showed similar response profiles (Fig. 3c-f), the SG and CC3 readouts showed notable differences with some drugs and not with others (Fig. 3c-h). Exposure to BORT generated no response by any measure. However, AAG, GELD, and MOC showed stronger CC3 responses than SG responses (Fig. 3c-h). This pattern may potentially reflect the time course of cell response with early activation of apoptosis, detected by CC3, preceding terminal cell death measured by SG. In contrast, PARTH and YM only showed strong SG responses, suggesting the possibility of non-apoptotic mechanisms of action for these drugs in our experiments (Fig. 3c-h). Thus the determination of responses depends on both the specific drug tested and the type of assay used as a readout. Finally, we compared dose-responses obtained for U87 cells *in vitro* to those for slices derived from U87 flank xenograft tumors. We found differential responses to particular drugs, even after accounting for the effects of different culture media used for *in vitro* and slice cultures (Fig. 3i). Most notably, while U87 was highly sensitive to BORT and MLN *in vitro*, no response was observed in slices, even at supramaximal doses (Fig. 3g). Together, these experiments demonstrate the critical impact that biological substrates and different assays can have on the interpretation of drug responses, such that both must be carefully considered when designing pre-clinical drug response platforms.

### Drug responses in intracranial derived xenograft slice cultures

U87 glioma flank tumor models proved useful for assay optimization and for characterization of slice cultures, but do not account for potential unique intracranial factors that could influence drug responses when grown in the brain, the native location of glioma tumors. Therefore to assess the potential modifying influence of the brain microenvironment, we tested drug sensitivities of slice cultures from intracranial (IC) xenograft tumors generated from U87 cells and GBM8 cells. We labeled the GBM8 cells with the red-fluorescent mCherry as GBM8 cells form invasive tumors. We first analyzed baseline proliferation, cell death, and cellular composition of IC slice cultures (Supplemental Fig. S6). Similar to U87 flank slices, U87 IC slices demonstrated a peak in cell death (SG and CC3) by 1-2 days, with a gradual but more variable decrease in proliferation (Ki-67) (Supplemental Fig. S6a,c,d,f,h,i). GBM8 IC slices showed a delayed increase in CC3 at 5 days, but a progressive growth from days 1-5 based on overall mCherry fluorescence (Supplemental Fig. S6e,g,j). Like with flank slices, we observed endothelial cells (CD31), immune cells (CD45), and macrophage/microglial cells (IBA-1) in U87 and GBM8 IC slices during the first few days in culture (Supplemental Fig. S6k). These data indicated that U87 flank and IC (U87 and GBM8) cultures experienced similar growth and stromal cell changes over time, except that GBM8 IC slices continued to grow for a longer time.

Direct comparison of U87 slices derived from flank vs IC tumors suggested that the unique microenvironments of these sites can influence selected drug responses. In slice cultures of U87 IC tumors, we assessed responses to BORT, PARTH and YM by mean SG fluorescence, SG index, and CC3+ area (Fig. 4a-d). When we compared the results to those obtained previously with U87 flank tumors (Fig. 3c-h), we found differences that depended on the tumor growth location. Compared to DMSO controls, BORT gave an ∼2 fold SG response (mean fluorescence only) in IC tumors, while no response was observed in flank tumors. Conversely, 100 µM PARTH gave only a small, statistically insignificant SG response in IC tumors, but a large SG response in flank tumors. YM, tested at 4 concentrations, showed similar SG responses (mean fluorescence and SG index) for both types of tumor slices, though starting at a slightly lower concentration in flank tumors (0.3 µM) than in intracranial tumors (1 µM). As with flank tumors, these three drugs did not produce CC3 responses in intracranial tumors. These differences suggest that the two microenvironments influence drug responses, although potential confounding factors (tumor size, cell proliferation, e.g.) may also contribute.

**Fig. 4:**
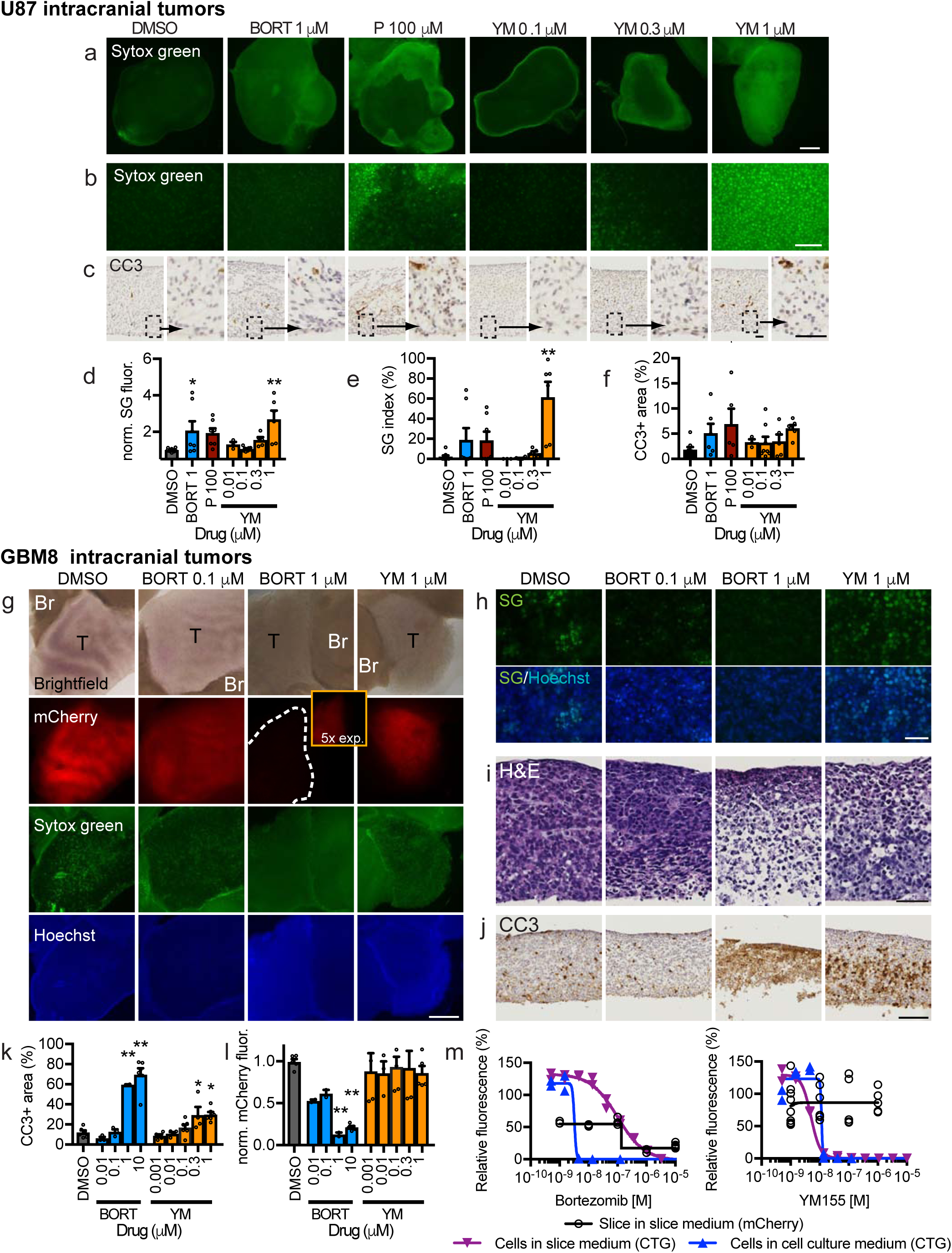
Intracranial xenograft drug response in slice culture. **a,f**) Cell death detected in U87 intracranial xenograft slices after 2 or 3 d treatment by SYTOX Green (SG) (**a,b**) or cleaved caspase 3 (CC3) staining (**C**). Drugs used were bortezomide (BORT, 3 d), parthenolide (P, 2 d), YM155 (YM 2d) and DMSO vehicle control (0.1%, 2 and 3 d). Cell death was quantified by SG fluorescence (**d,e**) or by CC3 staining (**f**). **G-H**) GBM8 glioma stem cell intracranial xenograft drug response in slice culture. **g**) mCherry-labeled GBM8 tumor (T) and adjacent brain (Br), with cell death changes detected by loss of mCherry, as well as by changes in SG/Hoechst (**g,h**) H&E (**i**), and CC3 (**j**) staining. Bortezomide reduced GBM8 mCherry signal (**k**), whereas both drugs increased % CC3-stained area (**l**). Dose-response curves for GBM8-mCherry xenograft slices versus the original GBM8 cells grown in glioma cell culture (blue line) or slice culture (purple line) medium. Cell death was assessed by CellTiter-Glo for GBM8 cells, and by loss of mCherry fluorescence for slices. Average ± SEM with n=8,7,7,3,8,5,6 slices per drug condition (**d-f**) except for n = 6 (BORT 1) and n = 5 (P 100) (**f**); n=6,3,3,3,3,6,5,4,6,5,6 slices per drug condition (**k,l**), and duplicates for cell lines (**m**). One-way ANOVA versus DMSO with Dunnett’s multiple comparison test. *p<0.05, **p<0.001. Scale bars: 1 mm (**a,g**), 100 µm (**b,j**), 25 µm (**c)**, 50 µm (**h,i**).

In slice cultures from IC tumors derived from the GBM8 stem cell line, we saw a different pattern of drug responses (Fig. 4g-m). The red fluorescent mCherry label in the GBM8 tumors allowed us to quantify drug responses as loss of mCherry fluorescence (normalized to DMSO control), an indicator of reduced viability. We also quantified increases in CC3+ area, as a measure of apoptosis, but did not quantify SG in these experiments because of high background signal with DMSO controls. We treated the GBM8 slices with two drugs, BORT and YM, at multiple concentrations. Representative images of brightfield, mCherry, SG, Hoechst, H&E and CC3 immunostaining are shown in Fig. 4g-j. We observed a robust loss of mCherry fluorescence when treated with 1 and 10 µM BORT, but no response to YM (Fig. 4g,k). GBM8 responses to BORT mirrored those of the GBM8 cells *in vitro* (CellTiter-Glo viability), but not to YM (Fig. 4m). The loss of mCherry signal with BORT was accompanied by marked increases in CC3+ areas at corresponding doses (Fig 4j,l), as well as by marked tissue disruption and nuclear fragmentation (Fig. 4h,i). For YM, an increased CC3+ area was seen despite no loss of mCherry fluorescence (Fig 4j,l). The detection of clear apoptosis with YM was not seen in U87 flank or intracranial tumors despite strong SG signal, even at a lower concentration, potentially reflecting a non-apoptotic mechanism in those cells. Note that the high background signal with DMSO controls (Fig. 4g,h) made SG analysis at low or high power problematic; the SG background with DMSO did not correlate with a loss of mCherry (Fig. 4g,k), histologic cell death, or significant CC3+ area (Fig. 4j,l). Together these data using two different GBM models (from U87 and GBM8 cell lines) demonstrated differential drug responses based on the culture platform (*in vitro* vs. slice) and the tumor location (flank vs. intracranial). Additionally, these results demonstrate how orthogonal assays (SG, CC3+ area, mCherry loss) can provide critical flexibility and sensitivity for drug response readouts.

### Dose-response readouts on device and comparison with off-device sensitivity

Next, we performed CP dose response curves on the microfluidic device using multiple slices of U87 flank xenograft cultures (Fig. 5). As shown in Fig. 5a, three slices placed next to each other were exposed to multiple CP concentrations on-device with intervening buffer lanes. At the end of the 2 day drug exposure, Hoechst was run in the drug lanes to mark the locations of drug delivery. We then removed the membrane and slices from the device and exposed the whole tissue to SG to evaluate cell death (Fig. 5a,c). Off-device control slices were treated with CP and SG in parallel (Fig. 5b,c). Significant SG cell death responses to the two highest concentrations of CP were seen on-device and off-device. Quantification of mean SG fluorescence (directly over the drug lanes on-device identified by Hoechst, as depicted in Fig. 5a) confirmed similar drug response patterns on and off the device, although the responses on-device were higher at the two highest doses (Fig. 5g). We also observed similar responses to the highest doses of CP, on- and off-device, detected by CC3 immunostaining of cross-sections (Fig. 5d-f). To identify drug lane locations, we imaged Hoechst in the same or adjacent cross-section while dry before hydration. Control experiments on-device with widely spaced delivery of high concentrations of CP showed cell death responses to CP (SG and CC3+) that extended, at the highest dose, from above the treated lanes, past the first adjacent buffer lanes, but not above the second lane over (Supplemental Fig. S7). These data clearly demonstrated the ability to assess dose responses of tumor slice cultures on the microfluidic device, and that on-device responses were similar to, or slightly more robust than, those obtained off the device.

**Fig. 5:**
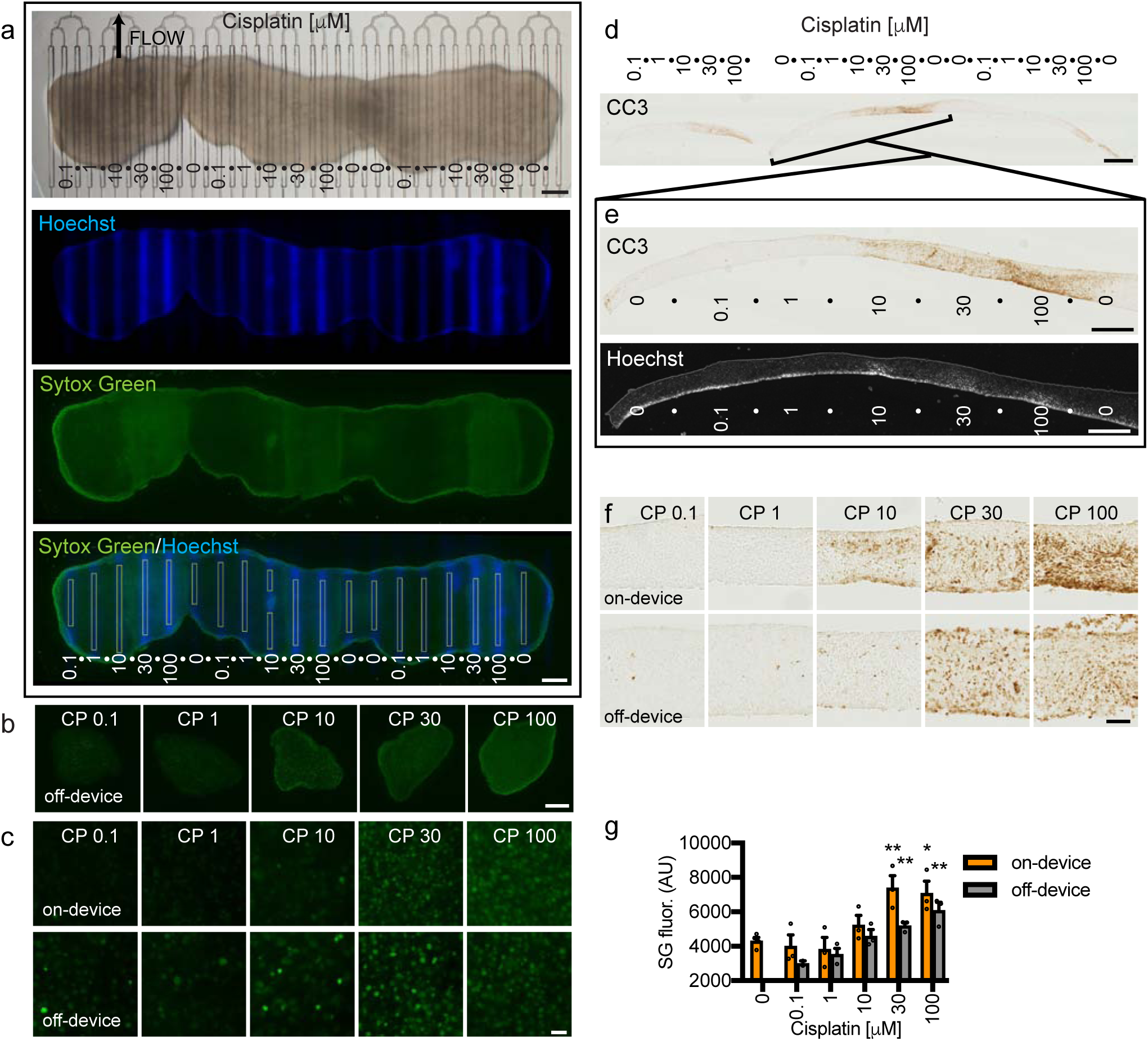
Xenograft slice culture dose-dependent drug response on and off the device. **a**) Three U87 flank glioma slices on a microfluidic device were treated with different doses of cisplatin (doses in µM) from days 1-3, followed by Hoechst dye in the drug lanes, then SYTOX Green (SG) staining of the entire undersurface. Drug lanes were separated by buffer lanes (•). SG mean fluorescence intensity (boxed lane regions, 100 µm wide) was used to quantify cell death (bottom panel). **b**) Low power images of SG staining from xenograft slices treated with drugs off-device in parallel. **c**) High power images of SG staining of slices treated on-device or off-device. **d,e**) Apoptotic cell death in cross-sections was detected by cleaved caspase 3 (CC3) staining. Lane locations were identified by alignment with Hoechst fluorescence. **f**) Higher resolution images of CC3 staining on-device or off-device. **g**) Cell death in response to CP on or off-device, measured by SG mean fluorescence. Average ± SEM with n=3 (lanes/slices) except for n=4 for 0 and n=2 for 0.1 µM CP tested off-device. One-way ANOVA versus 0.1 µM with Dunnett’s multiple comparison test separately for on and off-device. *p<0.05. **p<0.01. Scale bars: 1 mm (**a,b,d**), 100 µm (**c,f**), 500 µm (**e)**.

### Multiplexed drug testing on the microfluidic device

Next we performed multiplexed drug testing with the microfluidic device using four drugs on U87 flank xenograft slice cultures (Fig. 6). In the device experiments, each drug and the DMSO vehicle control were tested three times, each on a different slice on the same device. Hoechst marked the drug delivery lanes. SG, applied over the whole tissue, revealed cell death responses to CP, MOC, and PARTH, but not to BORT or to the DMSO control, as seen previously off-device (Fig. 3c-g). This response pattern was also visualized by an intensity profile across the tissue (Fig. 6d). Off-device controls run in parallel showed the same overall pattern of cell death for the drugs both at low power (Fig. 6b) and at high power (Fig. 6c). As quantified in Fig. 6i for mean SG fluorescence, each of the three repeats on-device showed strikingly similar levels of cell death (mean SG fluorescence), and the off-device controls showed a similar response pattern. An independently run second experiment showed the same results on and off-device (Fig. 6i). When we looked at apoptosis by CC3 immunostaining, we saw strong responses with CP and MOC, and a weak response with PARTH (Fig. 5e,g). This pattern was consistent with CC3 staining of off-device controls (Fig. 6h) as well as with the patterns seen off-device previously (Fig. 3). H&E staining also revealed histological changes for CP, MOC, and PARTH consistent with cell death (Fig. 6e,f). We also noted reproducible differences in the lateral extent of drug response. PARTH treatment resulted in narrower and shallower responses with SG and CC3 as compared to MOC and CP treatments (Fig. 6a,d,e,g). While this restriction may be due, in part, to differences in drug delivery, it may also be due to the steep dose response curve of PARTH (Fig. 3). These experiments established the potential utility of the device for multiplexed testing of drug panels on tumor tissue by demonstrating reliable and reproducible readouts for multiple drug responses between device runs and when compared with off-device responses.

**Fig. 6:**
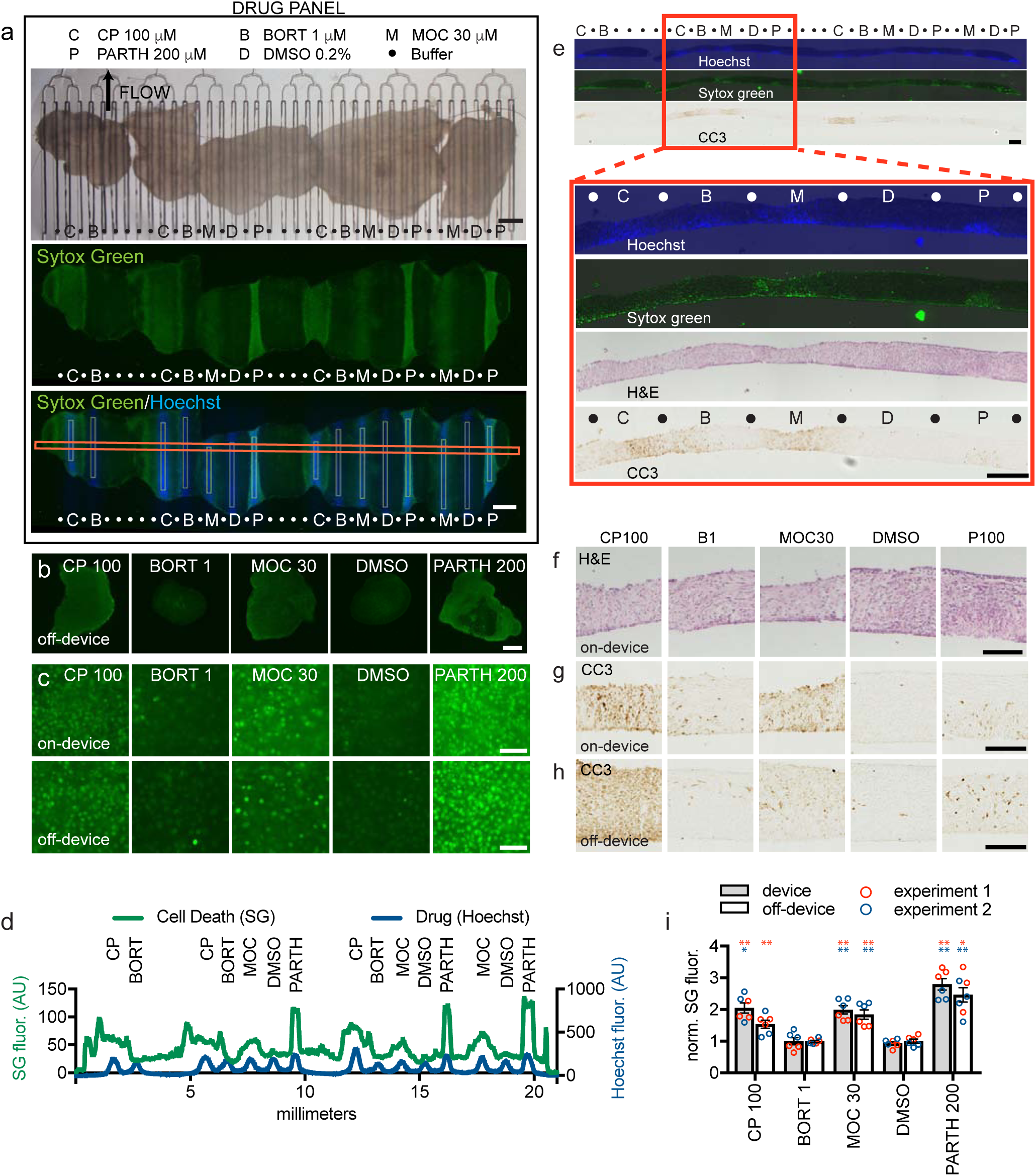
Multiplexed drug delivery to xenograft slices on the device. **a)** Five U87 flank xenograft slices on device treated with cisplatin (CP,C), bortezomib (B), mocetinostat (M), parthenolide (P), or DMSO (D) for 2 d beginning at d1. SYTOX Green (SG) staining, over drug lanes identified by Hoechst staining was used to quantify cell death in defined areas (lower panel, vertical white boxes 100 µm wide). The horizontal red box indicates the profile scan region for (**d**) below. **b)** SG staining shows similar drug responses in slices treated in parallel off-device (3 slices/condition). **c)** High-resolution images of SG responses on- and off-device. **d)** Horizontal profile across lanes (horizontal red box in (**a**)) identifies cell death (SG fluorescence) over drug lanes. **e-h)** Analysis of cross-sections reveals apoptotic cell death (CC3 staining) in drug lanes (markings as in (**a**)), with the boxed region showing aligned Hoechst, SG, CC3 and H&E images, also shown at higher magnification (**f-h**). **i**) Cell death as a function of treatment on- and off-device for 2 separate experiments with different devices and tissue. SG fluorescence (normalized to DMSO), shown with individual values (circles) for each lane/slice. Average ± SEM. n=3 each condition except for n=4 PARTH 200 off-device experiment 2. One-way ANOVA versus DMSO, Dunnett’s multiple comparison test, performed for on- and off-device and each experiment separately. *p<0.05. **p<0.01. Scale bars: 1 mm (**a,b**), 50 µm (**c**), 500 µm (**e)**, 200 µm (**f-h)**.

### Patient tumor slices on the microfluidic device

As a step towards clinical application, we then used the device to assess drug responses in human cancers using slices obtained from a glioblastoma (Fig. 7a-h) and a metastatic colorectal cancer (CRC) (Fig. 7i-n). For the GBM, we cultured three slices from the patient’s tumor overnight. The following day, we transferred the slices to the device and exposed them to CP for 24 hours (sometimes used for GBM chemotherapy alone or in combination with temozolomide), buffer (control for CP), staurosporine (STS, a non-selective protein kinase inhibitor), or DMSO (vehicle control for STS), as shown in Fig. 7a. Given the heterogeneity of the composition and baseline viability within patient tumors, we chose to use two different measures of apoptosis: CC3 immunostaining of sections as before, and CellEvent green-fluorescent staining of live, intact tissue, applied underneath the whole tissue immediately after drug exposure. We chose to use CellEvent for these experiments instead of SG because the higher signal-to-noise for CellEvent helped with the high tissue autofluorescence of the human GBM tissue. As seen in Fig. 7b, Hoechst marks the drug lanes, but the native autofluorescence of human brain tumors makes the CellEvent staining difficult to visualize in low power. High power images reveal blue Hoechst+ nuclei over drug lanes, and more dispersed CellEvent+ green dead nuclei over and next to the CP and STS lanes, as compared to their adjacent control drug lanes (Fig. 7c,d). Background subtraction of the high power images improved visualization and analysis. 4 of 5 CP repeats and 2 of 4 STS repeats showed obvious increases in apoptosis (Fig. 7e,f). When measured directly over the drug lanes, we could calculate a CellEvent index (% CellEvent+/total Hoechst+ nuclei) of ∼10% apoptosis for the CP lanes, and ∼8% for the two STS-responsive lanes (Fig. 7h). This heterogeneity in response was consistent with the patterns of apoptosis and tissue histology seen for CP and some STS lanes by CC3+ staining (Fig. 7g). Some baseline apoptotic signal was seen in the control and DMSO regions as well, consistent with similar and variable CC3 staining seen in day 0 tissue that had been fixed immediately (Fig. 7g). In this test of the device with a clinical GBM sample, we successfully tested 17 conditions on just three slices, a critical advantage when faced with a limited tissue sample.

**Fig. 7:**
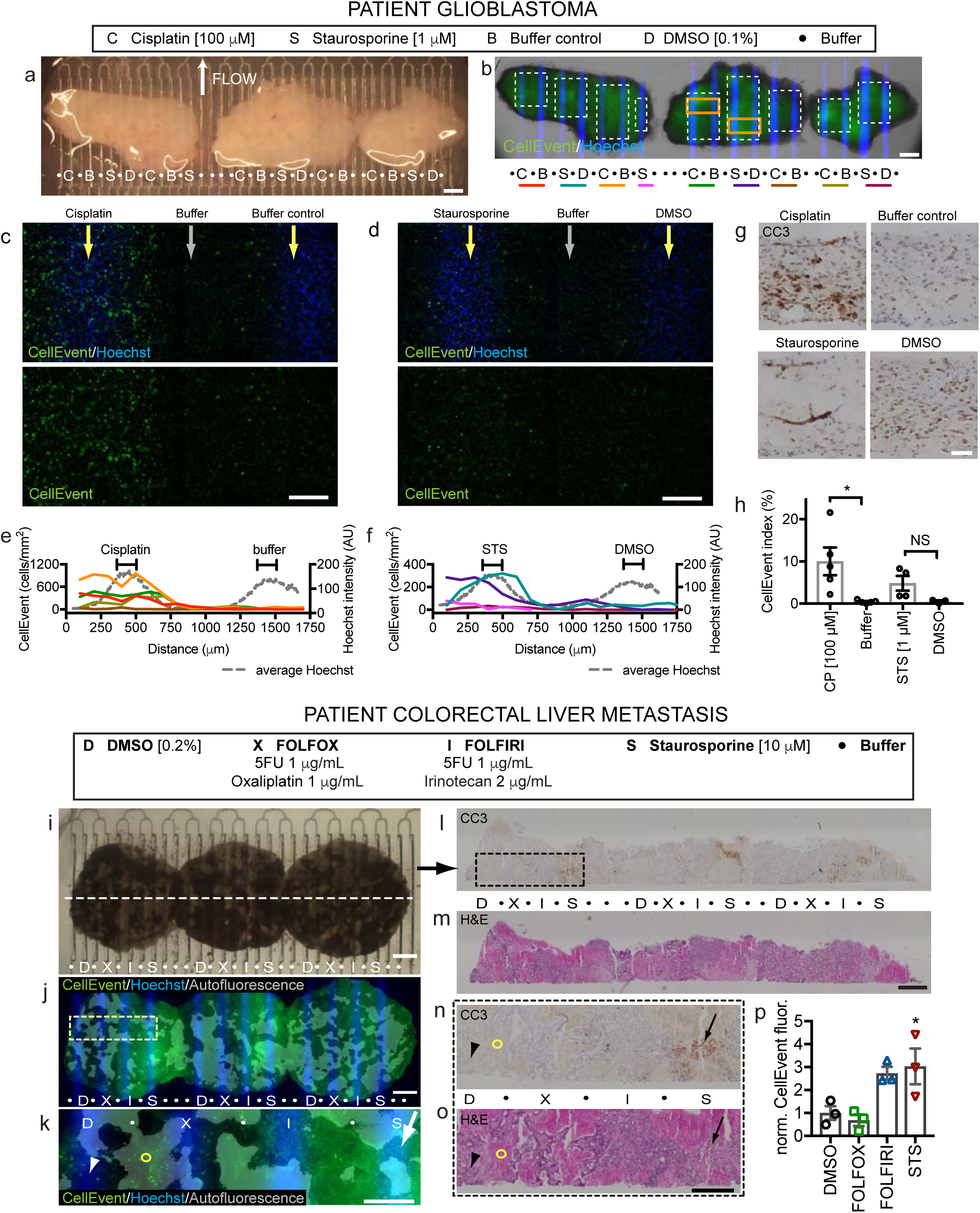
Patient glioblastoma and colorectal tumor drug responses on the device. **a-h**) Human glioblastoma resection slices treated for 24h on-device with cisplatin (C), staurosporine (S, STS), DMSO (D, control for STS) or buffer (B) starting at d1. **b**) CellEvent staining for apoptosis, shown at higher power for orange boxes in (**c,d**). **e,f**) CellEvent^+^ density (cells/mm^2^, bins) across lane pairs (same-color underlines and white boxes in (**b**)). Hoechst signal (gray lines) indicates drug delivery lanes. **g)** Cleaved-caspase 3 (CC3) staining detects apoptosis in sections. **h**) CellEvent^+^ cell fractions over drug lanes (100 µm wide). Average ± SEM, individual points overlaid, n=5,5,4,3. Student’s t-test for CP v Buffer and STS v DMSO, *p<=0.05. **i-n)** Colorectal cancer liver metastasis slices treated for 2 days on-device with FOLFOX (X), FOLFIRI (I), DMSO (D) or staurosporine (S). Hoechst identifies drug lanes. **j,l)** CellEvent shows apoptosis increased over drug lanes (arrow) versus DMSO (arrowhead). Red autofluorescence reveals old necrosis (circle). **l-p)** Sections stained for CC3 **(l)** or H&E **(m)**, boxed at higher magnification in **(n,o)**, show CC3 staining after staurosporine (red arrow) but not DMSO (arrowhead) treatment. **p)** Apoptosis quantified by CellEvent fluorescence normalized to DMSO (200 µm wide). Average ± SEM, n = 3. One-way ANOVA versus DMSO control, Dunnett’s multiple comparison test. *p<0.05. Scale bars: 1 mm (**a,b,i,j,l,m**), 200 µm (**c,d**), 500 µm (**e)**, 50 µm (**g)**, 500 µm (**k,n,o)**.

We also tested on device three slices of a patient’s colorectal carcinoma (CRC) liver metastasis with two standard chemotherapy regimens (FOLFOX and FOLFIRI, see Methods for the compositions of these mixtures) for 2 days (Fig. 7i-n). After the hypocellular stromal regions were removed from analysis (these regions have a strong red and green autofluorescence), the live apoptosis indicator, CellEvent, revealed a strong response to the staurosporine positive control, no response to FOLFOX or DMSO control, and a trend towards increased response to FOLFIRI (Fig. 6I-K, N). CC3 immunostaining confirmed apoptosis with staurosporine (Fig. 7l-p). Together these data demonstrated successful culture and drug testing of two different primary patient cancer slice cultures with our microfluidic drug testing platform.

## DISCUSSION

Rapid functional drug screens on intact patient tumor samples have the potential to improve both drug development and cancer precision medicine. Here we demonstrated proof-of-concept for a pre-clinical screening platform that incorporates microfluidic drug delivery to tumor-derived intact live tissue slice cultures. To establish feasibility, we developed a digitally-manufactured microfluidic drug delivery device, characterized therapeutically relevant biological changes of tumor slices in culture, and established protocols for drug response readouts both off- and on-device. We then successfully applied this approach to multiplexed drug response testing of both xenograft and human patient derived cancers.

The key findings of these experiments included: i) facile digital manufacturing of a 40 channel microfluidic device in PMMA plastic; ii) retention of cell proliferation, growth and stromal cell components in tumor derived slice cultures, iii) differential drug activity based on apoptotic vs generic cell death readouts, iv) the role of tumor microenvironment on drug responses evidenced by differential responses in 2D vs slice culture and flank versus intracranial derived slice cultures, v) successful use of the platform for quantitative multiplexed drug screening in xenograft and patient-derived slice cultures. Finally, these results were obtained within a week—a short enough time frame to inform clinical decision-making.

The design of our microfluidic drug delivery device permits straightforward testing of up to 20 single isolated drug conditions with both spatial and temporal control. One could test more drug conditions if adjacent channels contain drug, e.g. for different concentrations of the same drug for a dose response curve. This delivery platform is superior to our previously reported PDMS device in multiple ways^25^. Both platforms share user-friendly designs in multi-well plate format, with an open top and culture surface to permit easy placement of tissue samples, and flow controlled by a syringe pump connected to the single outlet. The PMMA device here drastically simplifies production by laser cutting using PMMA plastic instead of by multi-layer photolithography and alignment. The larger wells (∼5x) and wider fluidic delivery channels allow for longer run times with a reduced risk of clogging. Reproducible drug diffusion through tissue slices for up to 48 hrs can be achieved with drugs in every other lane for the PMMA device (20 conditions with 40 lanes), as opposed to every three lanes for the PDMS device (26 conditions with 80 lanes). The increased sampling area of the current device also allows a better assessment of intratumoral heterogeneity and variability in drug responses (Fig. 7, Supplemental Fig. S8). Other reported microfluidic platforms for drug testing have not achieved similar scales in testing of intact tissue; one group used micro-dissected tumors within up to 5 channels^12, 27^. Not in intact tissue, another group used small plugs of dissociated primary tumor cells to perform high-throughput testing of drug combinations, with brief culture and a single measure of cell death^34^. A final point is that the multiplexing capacity of our current device can be readily expanded: slice cultures on-platform can be turned 90 degrees to increase the number of discrete drug condition readouts from 20 to 400 (or 20 × 20) if the exposure window is expanded to a square^24^. Although these two perpendicular drug exposures would decrease the sampled area for any individual condition, it would substantially extend the search for potentially synergistic combinations and/or delivery protocols for both targeted and more conventional cytotoxic drugs^35, 36^.

A second important feature of our approach is the use of intact tumor slices that retain the 3D cellular and stromal architecture of the tumor microenvironment, potentially important determinants of drug responses^18, 37–39^. To the extent that tumor-derived slice cultures recapitulate the microenvironmental features of the parental tumor, the identification of situations in which the TME affects drug responsiveness in pre-clinical screening may be of practical importance. More in depth studies would be required to determine the mechanism and relative contributions of various features of the tumor microenvironment (cellular, structural matrix/soluble, and metabolic). For example, we observed distinct differences in drug response for proteosomal inhibitors in 2D vs. 3D slice platforms (Fig. 3). Interestingly, two prior studies have reported conflicting results with *in vivo* drug treatment of U87 xenograft tumors for one of those proteasome inhibitors, bortezomib, with one paper showing no effect^40^ and the other showing an effect^41^.

These experiments highlight the complexity in the comparison of slice culture drug studies to *in vivo* drug studies, tested over different time frames and with different criteria. An additional caveat of the xenograft slice cultures is that a human tumor grows in a non-native mouse microenvironment. Studies with tissue that retains the native microenvironment may also facilitate the testing of newer agents, such as shown in slice culture for immune check-point inhibitors^14, 42^ and adenovirus-based therapies^43^, alone or in combination with conventional targeted or genotoxic drugs. For example, Seo et al. recently demonstrated a response to immunotherapy with combined PD-1 and CXCR4 blockade in human pancreatic cancer slices^42^. The use of patient tumor tissue will permit studies with a native human microenvironment, and complementary studies in the future with syngeneic or transgenic mouse tumor models could exploit the intact mouse microenvironment. Thus, our microfluidic device should facilitate the multiplexed evaluation of combinations or gradients of immunotherapand traditional cancer drugs.

A third important general feature of our drug testing platform is the ability to use multiple response indicators to gain mechanistic insight into the extent and timing of drug responses. For example, we demonstrated the use of both apoptotic (CC3 and CellEvent) and generic (Sytox Green) readouts that provide a more robust snapshot of drug efficacy and likely mechanism of cell killing. Importantly, our approach is compatible with traditional pathology analyses that are used to evaluate cancer tissue in the clinic, such as the immunostaining shown here. Additional phenotypic or response readouts include measures of signaling pathways^44^; the induction of DNA damage responses (γH2AX staining)^8, 22^; cell proliferation (Ki-67 staining, Fig. S1); and cell migration and invasion^45^. All of these measures of intact tumor tissue response can be further extended by the analysis of slice culture tissue by biochemical^43, 44^, flow cytometric^32^, and genomic approaches^44^. It should eventually be possible to extend some of these imaging-based response indicators to real-time imaging readouts. This expansion of our platform would further improve our understanding of the time course and mechanisms governing therapeutic response in intact tissue slices and could enhance clinical relevance^46^.

Several potential limitations of our current microfluidic platform require further study or improvement to allow multiplexed drug profiling of a wide range of different tumor types. Tissue biophysical properties, and thus drug permeability, can vary within and between tumor types, especially in patient tumors. This variable will require additional tumor type-specific analyses to optimize drug delivery (directly with fluorescent dyes/drugs as in Fig. 2, or indirectly with drug effects for any drug as in Fig. S6), response readouts, and multiplexing. To help address the heterogeneity of tissue during experimental runs on-device, the internal control of adding dyes to drug delivery lanes during the experiment (here Hoechst dye for the last 2 h) provides one way to assess the variability of lateral diffusion, lane cross talk, and delivery. Drug responses must be normalized to the relevant treated tumor area, with careful consideration of best methods to evaluate the suitability of tumor/non-tumor regions for inclusion, and to ensure adequate sampling. Further refinement of our well-and-channel design should allow for slower continuous flow rates (currently ∼50 µL/hr/well), while maintaining effective drug delivery and lane separations. This improvement would decrease drug use while enabling longer on-device times that might be important for assessing responses to, e.g., immune-targeting drugs and biologics. Slice cultures capture many of the structural and cellular components of primary tumors and can be established for testing within days of tissue acquisition (see, e.g., Supplemental Figs. S2, S3). However, slices do not have an intact vascular or immune system. Thus it will be important to complement on-platform results in other, lower throughput, more physiologically intact, though nonhuman systems such as PDX models. Relative to the overall size of most patient tumors, our slice culture platform uses small tissue samples that, even when taken from different tumor regions, may not fully capture tumor genetic or metabolic heterogeneity (see, e.g., heterogeneous responses between different repeats from different regions of a GBM sample in Fig. 7c-f,h). These issues can be in part addressed by more systematic tumor sampling, including testing dispersed regions per drug and analyzing at a finer resolution. We could also modify and monitor other variables, such as hypoxia, that can alter therapeutic response^47^.

Our results provide proof-of-principle for a microfluidic platform-enabled, multiplexed drug profiling of human tissue slices. This platform is rapidly manufactured and it is intuitive to use, versatile, and compatible with a wide range of therapeutic agents including drugs, small molecules, ionizing radiation, and biologics. Moreover, depending on the intended use, several different types of mechanistically informative response indicators can be applied. Further refinement of this approach in the microfluidic drug delivery technology, response readouts, and biologic platforms should further enable the preclinical investigation of cancer biology and therapeutic responses in live tumor tissues. Future studies using PDX models, with differential drug sensitivities *in vivo* and reproducible tissue samples, represent a logical next step to evaluate the predictive power of our microfluidic slice approach. Eventually, clinical trials that incorporate concurrent testing of patient tissue samples with the device could establish the full clinical potential of this approach.

## METHODS

### Oncoslice device fabrication and operation

The Oncoslice device consists of a laser-cut PMMA 40-well plate whose central culture chamber (∼20 mm × 6 mm) has 40 microfluidic lanes driven by a single negative-pressure output. From top to bottom, the platform is an assembly of a PMMA 40 well plate (3/4 inch tall, laser cut with a ILS12.150D system) bonded with methylene chloride (Weld-On 4, Durham, USA) to a PMMA microfluidic chip. This chip layer consists of a 300 µm-thick channel layer thermally bonded, using chloroform and a heat press, to a 150 µm-thick PMMA bottom layer. The channel layer was cut using a VLS3.60 Universal Laser System. Microchannel dimensions were ∼120 µm wide and ∼80 µm maximum high for closed channels and ∼140 µm wide at the top and 300 µm high at the open area. The assembled microfluidic device was treated with oxygen plasma for hydrophilization and exposed to UV light for sterilization prior to use. Slice cultures grown on sterile hydrophilic PTFE membranes inserts (Millipore, 0.4 µm-diam. pores) were cut from the support and then directly placed with the membrane on top of the open lanes. The device was operated with a single negative pressure output at a flow rate of 2 mL/hr (average flow rate of 50 µL/h/lane) using an automated syringe pump. The device flow was interrupted briefly for ∼10 min as needed to empty the syringe.

### Fluorescent dye experiments

Using the device, slices were exposed to doxorubicin (DOX, 10 µM, Selleck) or Hoechst 33342 (16 µM, Invitrogen) for different time periods beginning on day 1 (48h) or day 2 (4h). Lanes were briefly rinsed with phosphate buffered saline (PBS, Invitrogen) for the last 10 minutes then the tissue was frozen and cryosectioned (10 µm for 48h and 20 µm for 4h). Fluorescent images were quantitated using the open-source image analysis program Fiji (https://fiji.sc/). For vertical profiles (50 µm wide), 2-4 adjacent or near adjacent sections were averaged for each location. For horizontal profiles (25 µm wide spaced 50 µm apart, starting at 25 µm above the surface), 3 near adjacent fluorescent images were analyzed. These images were aligned using an average of the lane centers determined for individual lanes by region of maximal fluorescence along a horizontal profile. Curved profiles were drawn following the shape of the membrane surface of the section.

### Cell culture and drug screening

U-87 MG (U87) cells (ATCC) were grown in DMEM/F12 (Invitrogen) supplemented with 10% fetal bovine serum (VWR) and penicillin/streptomycin (Invitrogen). GBM8 cells^48^ were grown in Neurobasal medium supplemented with B27, N2, Glutamax, penicillin/streptomycin (Invitrogen), as well as with growth factors (EGF 20 ng/mL and FGF 20 ng/mL, Preprotech). Both cell lines tested negative for *Mycoplasma* and their identity was confirmed by microsatellite analysis. For slice medium experiments, cells were grown in Neurobasal-A (Invitrogen) with 25% heat-inactivated horse serum (Sigma), Glutamax (Invitrogen), and 2× penicillin/streptomycin (Invitrogen). U87-EGFP cells were created by infection with lentivirus made from pLL3.7 (Addgene #11795). The drug screens were performed by the Quellos High Throughput Screening Core (University of Washington, Seattle) with CellTiter-Glo (Promega), as well as with CellTox Green (Promega). Drug treatment began on day 1 and was performed in duplicate (10-point, 3-fold dilutions from 10 µM for the primary screen, or from 10-100 µM for the secondary screen). Total fluorescence was read with a plate reader on days 2, 3, and 4 for CellTox Green (secondary screens only, added on day 1), and on day 4 for CellTiter-Glo. The signal was normalized to the signal from the DMSO vehicle control.

### Xenograft mouse model

Mice were handled in accordance with a protocol approved by the University of Washington Animal Care and Use Committee. Male athymic nude mice (Taconic, Foxn1^nu^) aged 4-10 weeks were injected either intracranially (100,000 cells at the dorsal edge of the right striatum) or subcutaneously in the flank (0.5-1 million cells in 200 µL of serum and antibiotic free medium). Mice with orthotopic tumors were sacrificed once they demonstrated signs of morbidity (2-4 weeks). Mice with flank tumors were sacrificed before tumor volume reached 2 cm^2^ (2-4 weeks).

### Human tissue

Human tissue was obtained with written informed consent and treated in accordance with Institutional Review Board approved protocols at the University of Washington, Seattle. A biopsy from a 52-year-old male with GBM was embedded in 2% low melt agarose (ISC Bioexpress) in PBS for sectioning. A liver metastasis biopsy was obtained from a 53-year-old female with metastatic colon cancer post multiple treatments and immediately placed in Belzer-UW cold storage medium (Bridge-to-Life Ltd).

### Slice culture

250 µm-thick brain tumor slices (except 300 µm for human GBM tissue) were cut with a 5100mz vibratome (Lafayette Instrument) and cultured on top of PFTE 0.4 µm transwell membranes (Millipore) in 6-well plates. Brain slices were prepared in ice-cold, Gey’s balanced salt solution (Sigma, St. Louis, MO) bubbled with carbogen (5% CO2 and 95% O2). The culture medium underneath was Neurobasal-A medium (Invitrogen) with 25% heat-inactivated horse serum (Sigma), Glutamax (Invitrogen), 2× penicillin/streptomycin (Invitrogen), and growth factors (EGF 20 ng/mL and FGF 20 ng/mL, Preprotech or Invitrogen). The medium was changed three times per week. Drugs were added to medium from day 1-3 unless otherwise specified. Slices from at least two animals were analyzed for each drug except for geldanamycin (one animal). For CRC slices (obtained from H. Kenerson and R. Yeung, University of Washington, Seattle), 250 µm-thick slices were cut with a Leica VT 1200S vibrating microtome, then cultured on the PFTE inserts with shaking. Culture medium was Williams’ Media E (Sigma) supplemented with nicotinamide (12 mmol/L), L-ascorbic acid 2-phosphate (50 mg/ml), D-(+)-glucose (5 mg/ml) from Sigma; sodium bicarbonate (2.5%), HEPES (20 mmol/L), sodium pyruvate (1 mM), Glutamax (1%), and penicillin-streptomycin (0.4%) from Gibco; and ITS+ Premix (1%) and human EGF (20 ng/ml) from BD Biosciences.

For off-device experiments, one to two hours before the end of the experiment, nuclear dyes were added to the growth medium: Hoechst (16 µM) and SYTOX Green (SG, Invitrogen 0.1 µM). Then slices were washed three times with PBS for 5 minutes at room temperature. Slices were fixed with 4% paraformaldehyde overnight, then cryoprotected with two changes of 30% sucrose/PBS. Low and high power images of the bottom surface were taken with the slices on the membrane. For on-device experiments, Hoechst (16 µM) was added to the drug lanes 2 hours before the end of the experiment. At the end of the treatment period, the slice and membrane were placed onto a new cell insert. After 3 washes with PBS, the slice was incubated with SG (0.5 µM) or CellEvent (Invitrogen, 1/1,000) in medium for one hour, washed 3 times with PBS, then fixed and processed as above for off-device slices. For the GBM experiments, ethidium homodimer-1 staining for dead nuclei (red, 2 µM, Invitrogen) was also performed but the results were inconclusive.

Hoechst dye in dry cross-sectional cryosections stains nuclei on the bottom surface and thus identifies the membrane side as well as lane location for device experiments. Unfortunately, upon hydration, SG and Hoechst staining spreads to all nuclei, likely because the dyes cannot be completely cleared from the live tissue post staining and all cells are dead at that point.

### Immunostaining and image analysis

Tissue slices were cut in half with one half processed for paraffin sectioning (4 microns) and H&E. The other half was processed for cryosectioning (14 µm thick except for 10 µm for CRC) and immunostaining. For immunostaining, tissue was incubated for at least 30 minutes in blocking solution (TNB, Perkin Elmer, with 0.1% Triton X-100) and if a mouse primary, 30 minutes with Rodent Block M (Biocare Medical). Tissue was incubated with primary antibody (diluted in TNB) overnight at 4C. Primary antibodies were as follows: rabbit anti-Ki-67 (1/300 fluorescence 1/4,000 DAB), AbCAM, ab15580), rabbit anti-CD31 (1/60, AbCAM ab28364), rabbit anti-Iba-1 (1/1,000, WAKO), rabbit anti-active caspase 3 (Asp175) (1/600, Cell Signaling, Antibody #9661), mouse anti-HuNu (1/500, AbCAM ab191181), goat anti-vimentin (1/300, Chemicon AB1620), rabbit anti-CD45 (1/1000, AbCAM, ab10558). For immunofluorescence, tissue was incubated with donkey secondary antibodies for 1 hour (Invitrogen, 1/500 diluted in TNB + 0.1% Triton X-100) then coverslipped with Fluoromount with DAPI (Southern Biotech). Images were taken using a Nikon TiE inverted microscope and NIS Elements software. For peroxidase staining, tissue was pretreated with 0.6% hydrogen peroxide in methanol for 30 minutes. For human tissue, antigen retrieval was performed for 30 minutes in 10 mM sodium citrate, 0.05% Tween 20 (Sigma), pH 6.0. After primary antibody treatment, tissue was incubated with anti-rabbit peroxidase polymers (Vector Labs MP7401) for 30 minutes, then with DAB solution (Vector Labs) and lightly counterstained with hematoxylin. Stained slides were scanned with a Hamamatsu Nanozoomer then DAB staining was quantified using Visiopharm software by the Histology and Imaging Core at the University of Washington. Confocal images were taken with a Nikon A1R confocal provided by the Garvey Cell Imaging Lab (University of Washington).

SG and CellEvent image analysis on tiled 2× images taken of intact slices was performed using Fiji. A central region was selected for each slice using the Hoechst channel avoiding approximately 1 mm of the edge. Five circular regions outside the slices were selected for background. The background was calculated by averaging all values for that experiment for SG. After background subtraction, the total average SG fluorescence was calculated for each region relative to that of DMSO (for drug treatments) or to that at day 3 (culture time course).

For 20× SG image analysis, unbiased image collection was performed by placing a 1 mm × 1 mm grid over the 2× images and acquiring 20× images in the centers of the boxes in which the tissue filled the box. A custom routine made on the open-source image analysis program, CellProfiler (Broad Institute, www.cellprofiler.org), was created to identify all nuclei in the Hoechst channel then measure the SG fluorescence after 3 part Otsu threshold correction of each image. A single threshold for positive nuclei was determined for each experimental set using the help of positive control crush lesions made at the time of nuclear dye labeling. For GBM8-mCherry fluorescence, a custom CellProfiler routine outlined the area of mCherry fluorescence and measured the mean fluorescence. The appropriateness of the selected area was confirmed by eye for all channels.

CellEvent image analysis was performed on images taken of live tissue just after viability staining. For the GBM experiment, the images were tiled 10× images covering each lane (as defined by Hoechst staining), 200 µm wide, and excluding the edges of the tissue. Initial background subtraction using a rolling ball radius of 25 pixels was performed with Fiji. Using CellProfiler, Hoechst and CellEvent images had nuclei counted using a three-part Otsu threshold. The threshold for all CellEvent images was determined by using the average value of the threshold calculated for all images that contained many clearly labeled cells. For the CRC experiment, CellEvent fluorescence was measured with CellProfiler on images covering each lane (200 µm wide and excluding the edges of the tissue). Mean fluorescence was measured after masking red autofluorescent regions using a single threshold determined over the entire tiled 4x image.

Immunofluorescence quantification was as follows. For Ki-67 analysis, a custom CellProfiler routine counted the total number of DAPI+ and of Ki-67+ nuclei in each image, which covered each full slice cross-section with tiled 10× images. The Ki-67 threshold was adjusted and kept constant for each experimental set. IBA-1 counts were performed blinded on 20× images and the total number of DAPI+ nuclei was calculated using the same Cell Profiler algorithm used for DAPI+ nuclei above in the Ki-67 experiments. Quantification of GFP/HuNu/DAPI/vimentin staining of confocal images (20×) was performed blinded by one individual and confirmed by a second individual for a small subset. Images were visualized with the help of Fiji. For vimentin counts, the total number of DAPI+ nuclei was calculated with the ICTN plug-in.

## Materials

Drugs were diluted from DMSO stocks (10-200 mM), purchased from MedChem Express, except for temozolomide (TCI Chemicals), doxorubicin (Selleck), and cisplatin (MedChemExpress, 3M stock in dH20).

## Statistical analysis

Unless otherwise indicated, data were analyzed using either using a one-way ANOVA versus control with Dunnett’s multiple comparison test (drug experiments), or with a one-way ANOVA with Tukey’s multiple comparison test (culture experiments).

## Data Availability

The datasets generated and analysed during this current study are available from the corresponding authors on reasonable request.

## Acknowledgments

We would like to thank Heidi Kenerson and Ray Yeung for the CRC cancer tissue slices, as well as Max Rumaner, Christine Tan, Ryan Palermo for help with cell counts. We are indebted to Paul Yager’s group for training in and use of their laser cutting facility. Thanks to the University of Washington Pathology Core for lending equipment. We also thank John Silber, Daniel Silbergeld, and the brain tumor bank for providing the GBM tissue. This work was supported by a grant from the National Cancer Institute R01 CA181445-01A1 and an International Scholars award from the Consejo Nacional de Ciencia y Tecnología of Mexico for A.R.

## Author contributions

Contributions were as follows. L.F.H., A.R., K.C., A.M., R.J.M., R.C.R., and A.F. designed experiments. L.F.H. A.R., and K.C. performed experiments. L.F.H., R.L., and Z.D. performed analysis. L.F.H., Z.D., R.J.M., R.C.R., and A.F. wrote the manuscript.

## Competing interests

The microfluidic device is covered under US patent 20140113838A1, on which L.F.H., R.J.M., A.F., and R.C.R. are inventors. L.F.H. and A.F. are founders of OncoFluidics, LLC.

## Materials and Correspondence

Correspondence and materials requests should be addressed to Robert Rostomily at or to Albert Folch at.

---

**Supplemental Fig. S1.**
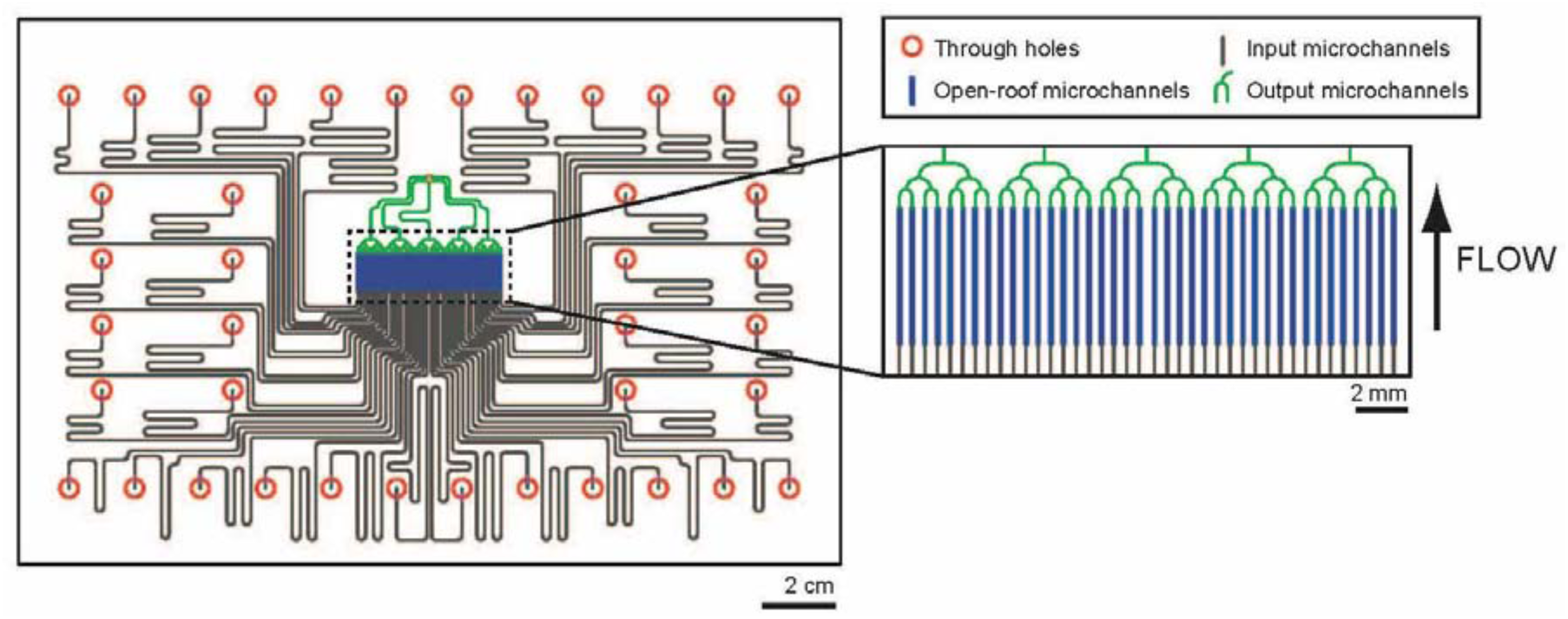
**Layout of device microchannel layer**. CAD drawings show the device microchannel layer, and the drug delivery area at higher magnification. Through holes (red circles) connect via input microchannels (grey) to the open-roof microchannels (blue), which will be covered and closed by the culture membrane and overlying tissue during device operation. Output microchannels (green) then combine into a single output.

**Supplemental Fig. S2.**
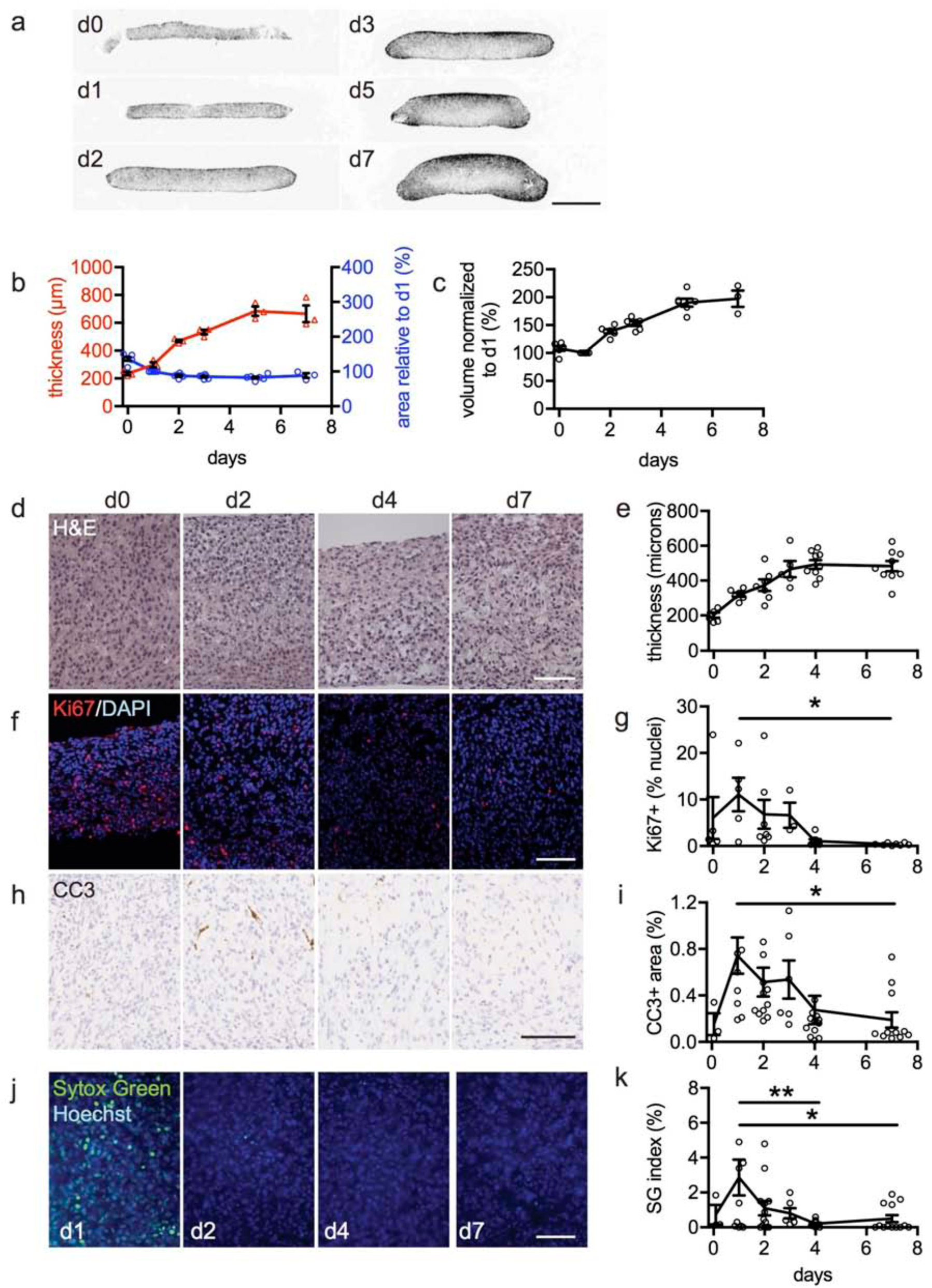
Characterization of cell growth, death, and proliferation in glioma xenograft slice cultures. U87 flank xenograft slices were cultured for up to 7 days, following by sectioning and staining to quantify slice appearance and cell growth/death. **a-c**) Low-power cross-sections of U87-GFP xenografts (membrane surface down) (**a**) were used to quantify changes in thickness and relative area (**b**) and in volume (estimated by thickness × area) (**c**) over time (n=3 each timepoint). **d-k**) U87 xenograft slices were used to assess overall histological appearance by H&E (D); to quantify changes in slice thickness (**e**, n=6-9 each timepoint); to measure cell proliferation by Ki-67-stained nuclear fraction (**f,g**; n=3-8); and to measure cell death by cleaved caspase 3 (apoptotic cell death; **h,i**; n=3-13) or SYTOX Green versus Hoechst staining (**j**,**k**; n=3-13). Both cell proliferation and cell death peaked on day 1, then declined over the subsequent 3 days in conjunction with a plateau in slice thickness. All curves represent the average ± SEM, one-way ANOVA, Tukey’s multiple comparison test, with * p<0.05, **p<0.01. Scale bar = 500 µm (**a**), 100 µm (**d,f,h,j**).

**Supplemental Fig. S3.**
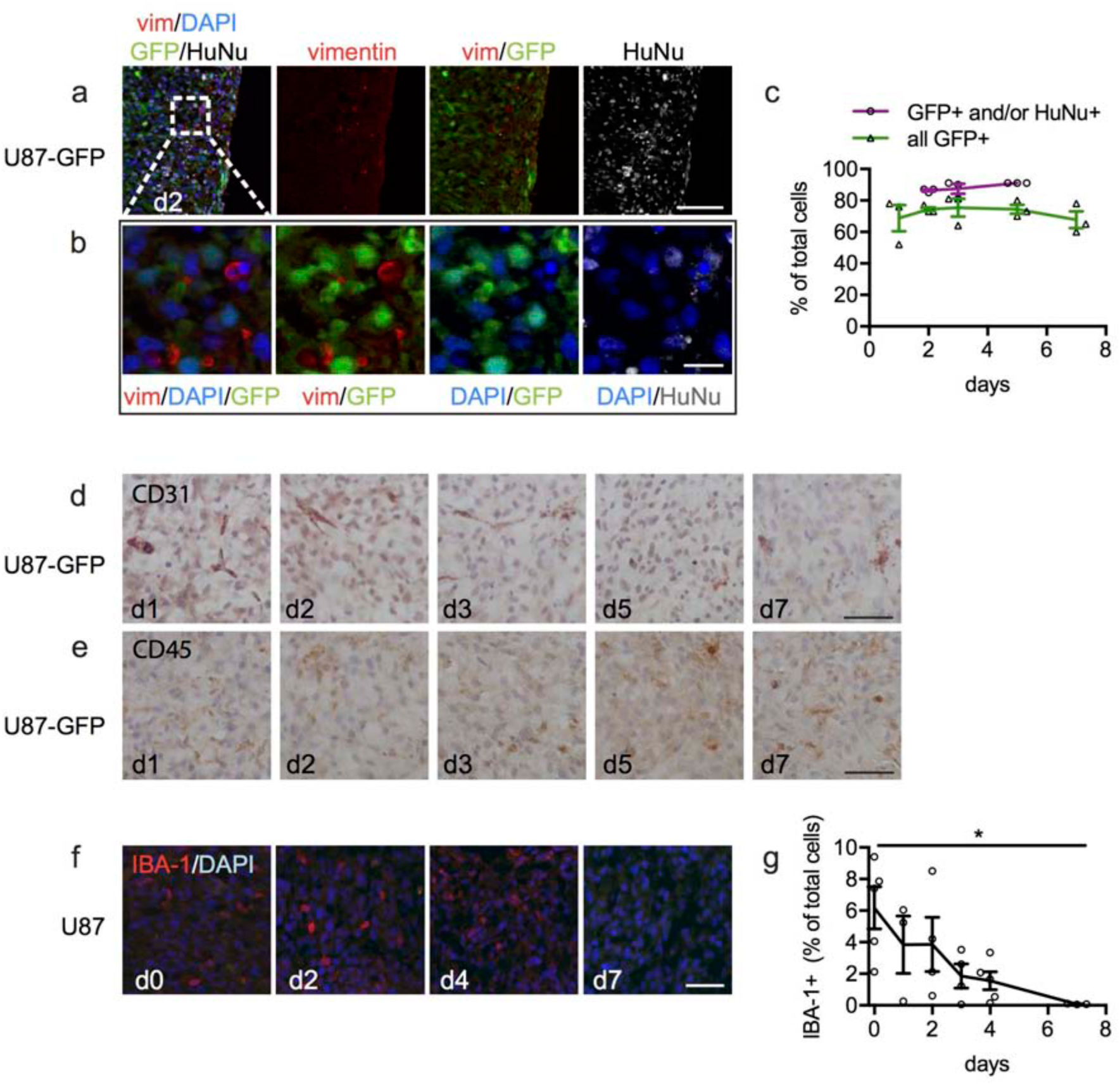
Xenograft cellular microenvironment. **a,b**) U87-GFP flank tumor slices were stained after different times in culture with antibodies to human nuclear antigen (HuNu), then confocal-imaged to identify tumor cells (GFP+ and/or HuNu+). The percentage of tumor cells was quantitated as a percentage of total cells (DAPI+) in (**c**) (ave ± SEM, n=3). Fibroblasts were identified in the same sections by antibodies to vimentin to identify mesenchymal stromal cells (vimentin+/GFP-/HuNu-). DAB and hematoxylin immunohistochemistry of the same U87-GFP slices revealed a gradual loss of CD31+ endothelial cells (**d**) and persistence of CD45+ immune cells (**e**). **f,g**) Antibody staining of U87 flank xenograft slices show a decline in IBA-1+ macrophage over 7 days in culture. Average ± SEM, n = 5,3,4,4,5,3. One-way ANOVA, Tukey’s multiple comparison test. * p<0.05 versus d0. Scale bar = 100 µm (**a**), 20 µm (**b**), 50 µm (**d-f**).

**Fig. S4.**
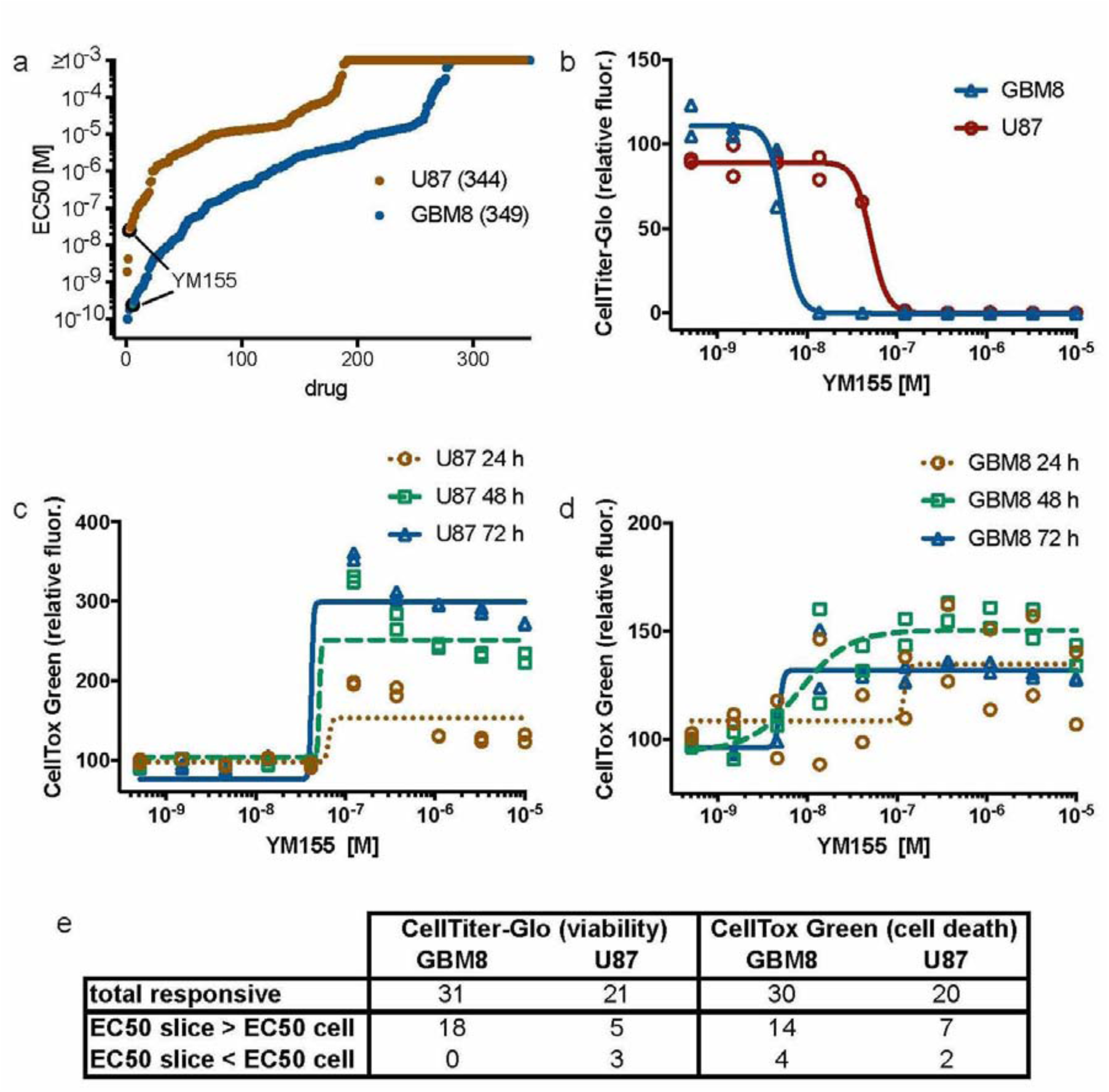
High-throughput drug screening of U87 glioma and GBM8 glioma stem cell lines. **A**) Waterfall plots of EC50 values from a primary screen of U87 and GBM8 cells with Selleck anti-cancer library drugs, with reduced viability at 72 h assessed by CellTiter-Glo. The total number of drugs screened (in parentheses) and the position of YM155 are indicated. **B-D**) Example from a secondary screen of U87 and GBM8 cells with 32 drugs, with YM155 dose-dependent growth suppression seen by CellTiter-Glo at 72 h (**B**), as well as a time course of YM155 dose-dependent killing of U87 and GBM8 cells seen by increased CellTox Green fluorescence at 24, 48, and 72 h (**C,D**). **E**) Summary statistics for the ability of the 32 drugs to suppress the growth or viability of U87 and GBM8 cells as a function of culture medium. Cell viability was quantified by CellTiter-Glo and cell death by CellTox Green for each cell type, grown in their routine culture medium (‘cell’) versus in slice culture medium. A ≥4.5 fold increase or decrease in EC50 was used as a cutoff and the number of drugs meeting each condition is specified. Each screen was performed in duplicate.

**Supplemental Fig. S5.**
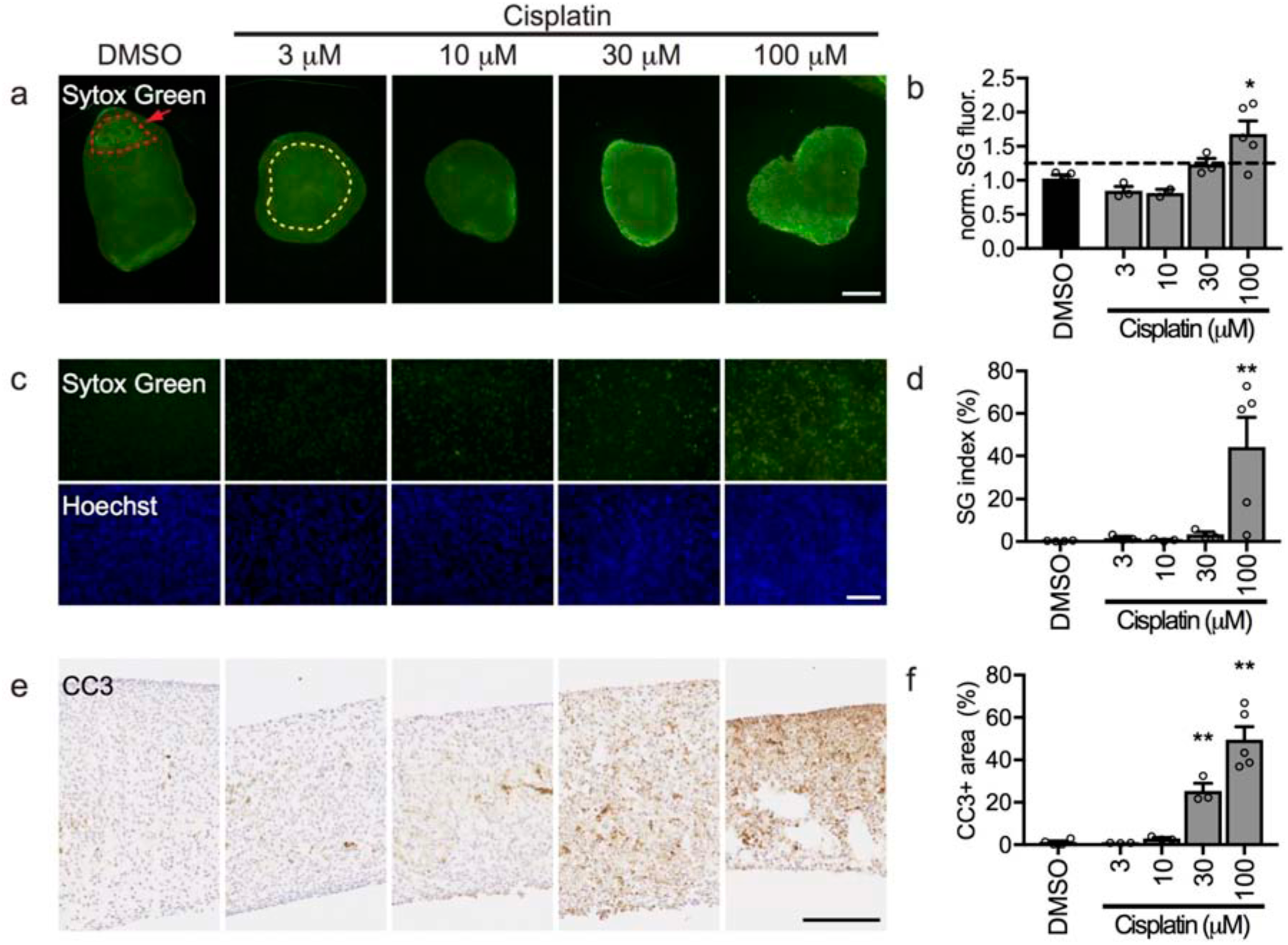
Dose-dependent increase in cell death after cisplatin treatment of U87 slices in culture. U87 flank tumor slices treated with different doses of cisplatin (CP) from days 1-3 were then exposed to Sytox Green (SG, green dead nuclear stain) and Hoechst (blue nuclear stain) before washing and fixation. Non-specific cell death was assessed in low power images (**a**) as overall fluorescence over the central area (dotted yellow line). A positive control crush lesion on the DMSO slice is outlined in red. Treatment with 100 µM CP led to increased cell death, measured as SG fluorescence normalized to DMSO control (**b**). (**c**,**d**) Analysis of high power images showed a similar increase in cell death at 100 µM CP, calculated as an SG index (% SG+ nuclei/all Hoechst+ nuclei). **e,f**) Apoptotic cell death was assessed by CC3+ immunostaining of cross-sections (**e**) and quantified as CC3+ area (**f**), revealing apoptotic cell death at 30 µM as well as at 100 µM CP. Average ± SEM; n=4,3,3,3,5 (**b,d,f)** except n=2 for 10 µM in (**b**). One-way ANOVA versus DMSO with Dunnett’s multiple comparison test. *p<0.05, **p<0.01. Scale bar = 1 mm (**a**), 100 µm (**c**,**e**).

**Supplemental Fig. S6.**
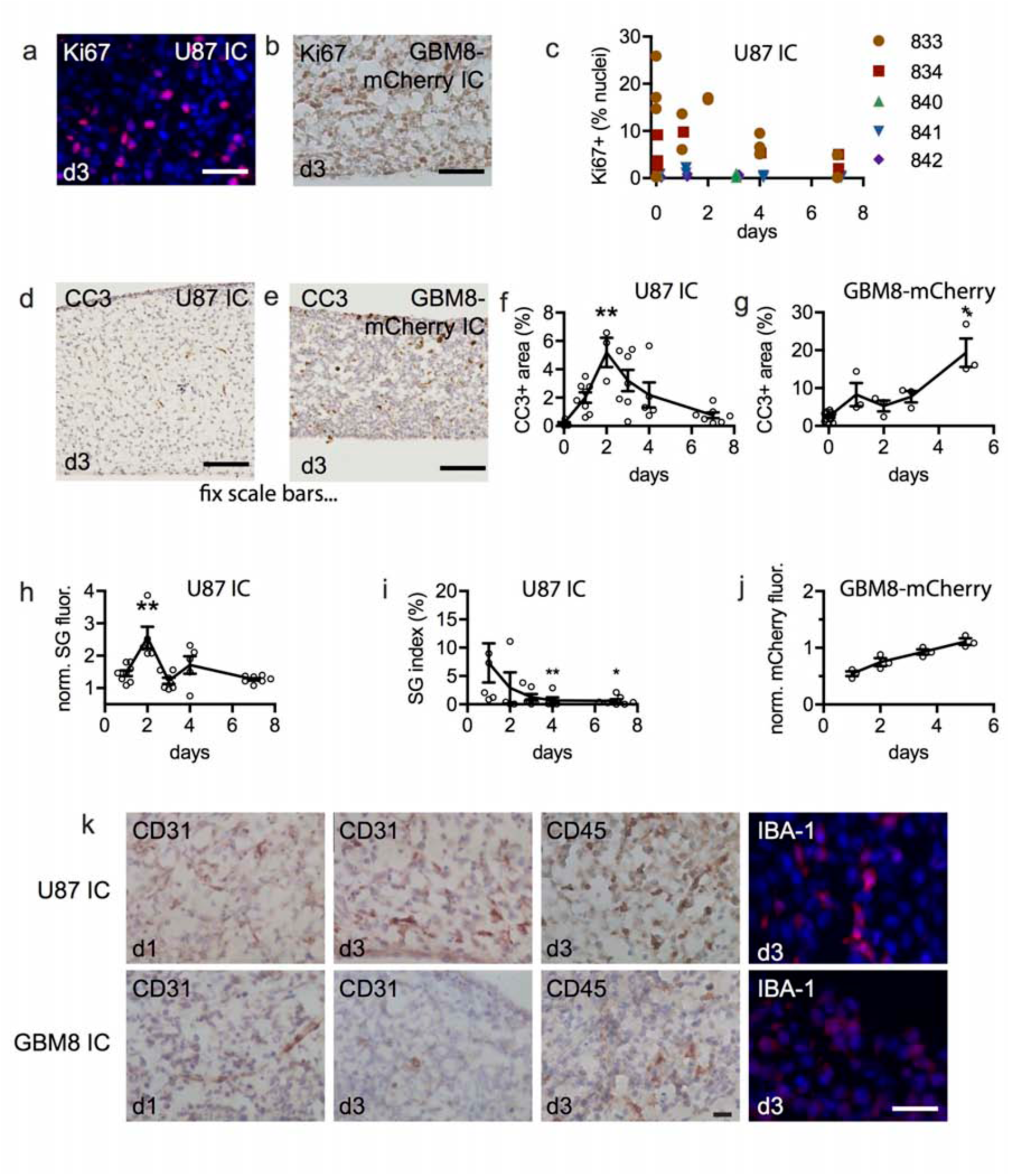
Characterization of intracranial xenograft slice cultures derived from U87 or GBM8 cells. **a-c)** Cell proliferation was detected in intracranial xenograft slices by Ki-67/Hoechst (**a**) or Ki-67/ hematoxylin (**b**) staining at d3. **c)** Time course Ki-67 staining as % of total nuclei, shown for individual slices from 5 separate U87-derived xenograft animals (number ID on right). **d-g)** Apoptotic cell death detected by cleaved caspase 3 (CC3) immunostaining in U87 over 7d, or in GBM8-mCherry xenografts over 5d, quantified by measuring CC3+ stained slice area (f,g). Overall cell death was quantified for U87-initiated xenografts by mean SYTOX Green (SG) fluorescence normalized to d3 (**h**), or by an SG index (% SG+ nuclei) (**i**). **j**) mCherry fluorescence (normalized to DMSO controls at d3 run in parallel) in GBM8-mCherry did not decrease over 5d. N = 3 per timepoint. **k**) Presence of additional cell types in U87 and GBM8 slice cultures shown by immunostaining for endothelial cells (CD31), immune cells (CD45), and microglia (IBA-1) at d1 or d3. Graphs show average ± SEM. N=6,8,3,7,5,7 (**f**); n=8,5,7,5,7 (**h**); n=6,4,7,5,7 (**i**); and n=3 for GBM8-mCherry (**g**,j) except for n=14 at d0. One-way ANOVA with Tukey’s multiple comparison test. *p<0.05, **p<0.01 compared to initial timepoint. Scale bar = 50 µm (**a**,**b**), 100 µm (**d**,**e**), 20 µm (**k**).

**Supplemental Fig. S7.**
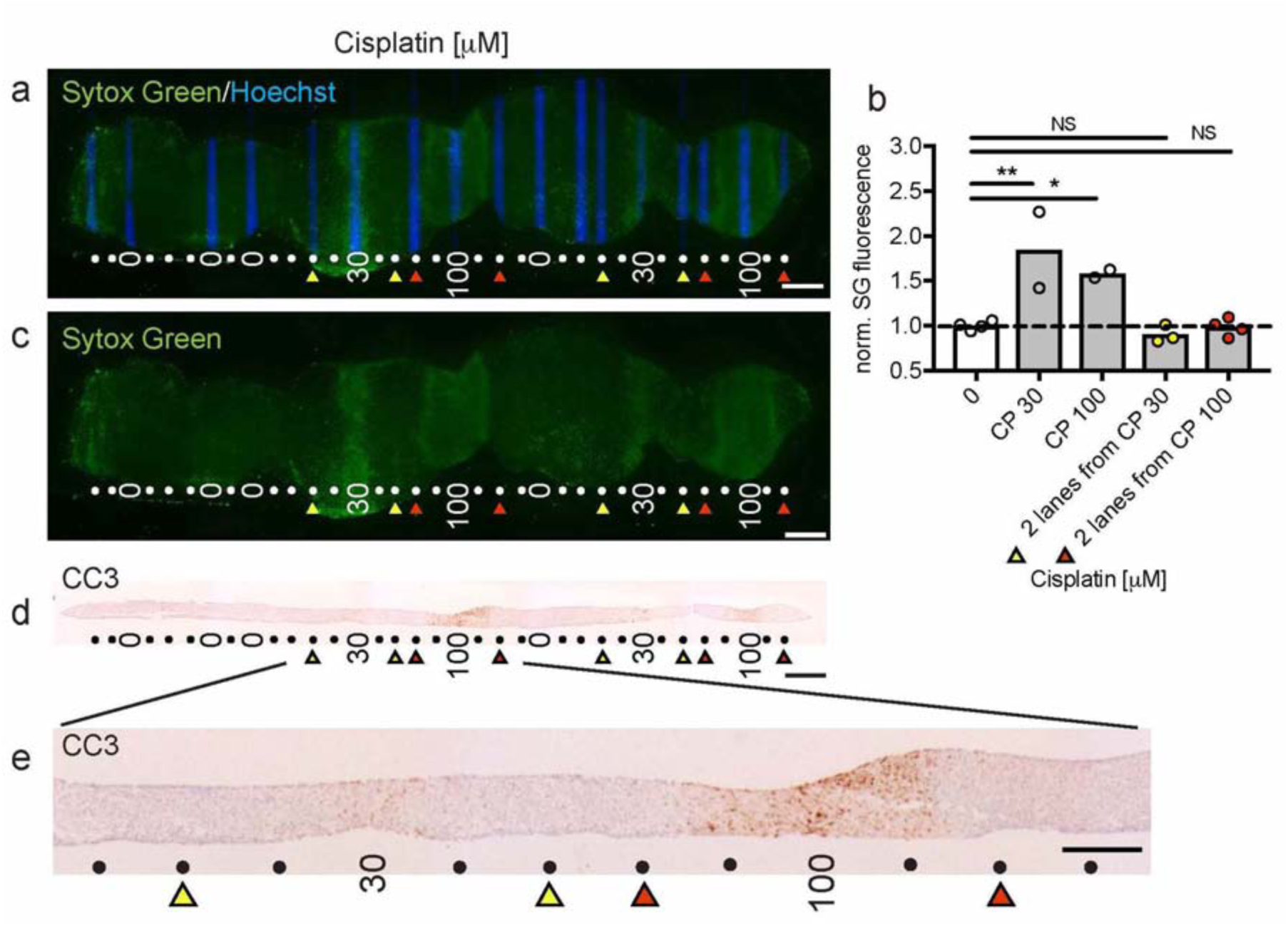
Restricted lateral spread of cisplatin-mediated cell death in slice cultures on the device. **a,c**) U87 flank xenograft slices were treated with different cisplatin doses (indicated in µM) for 2 d on-device after 1 d off-device, followed by staining with Hoechst dye delivered through drug lanes and SYTOX Green (SG) staining over the entire slice. Yellow and red triangles identify lanes located 2 lanes over from drug delivery lanes. **b**) Quantification of cell death by SG fluorescence, normalized to 0 (buffer alone), reveals a significant increase in cell death over drug delivery lanes, but not in lanes located 2 lanes away (identified by triangles as in panels **a** and **c**). **d,e**) Apoptotic cell death, detected by cleaved caspase 3 (CC3) immunostaining, was limited largely to the drug delivery lane with only modest lateral spread at the highest concentration tested (100 µM) that doesn’t reach 2 lanes away. The graph in (**b**) shows average and individual points. One-way ANOVA versus buffer control, Dunnett’s multiple comparison test. *p≤0.05. **p≤0.01. Individual points and average. N=7,3,4,5,5. Scale bar = 1 mm (**a,c,d**), 500 µm (**e**).

**Supplemental Table S1.**
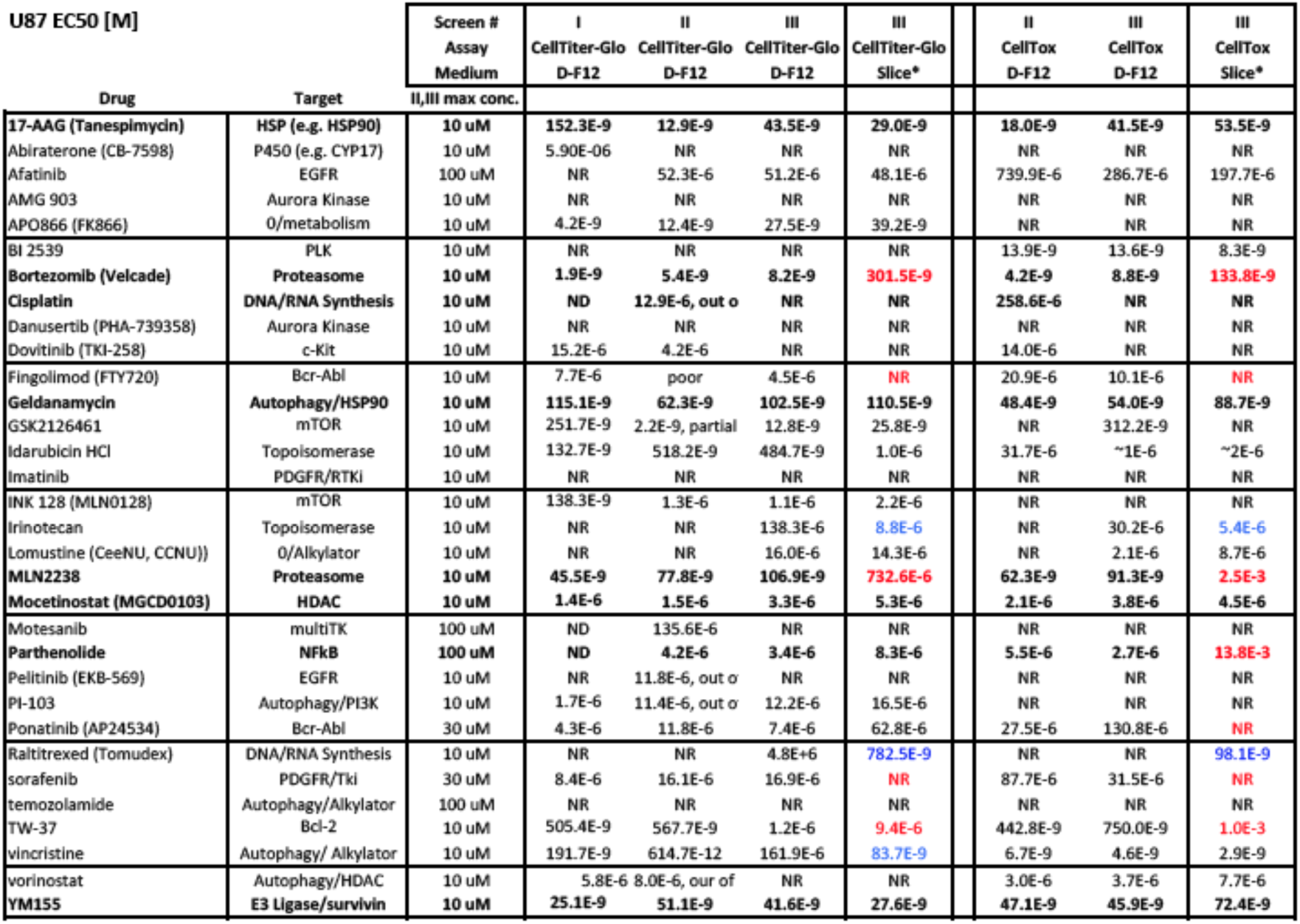
Secondary screen results with U87. 3 screen runs (I, II, and III) were performed with U87 cells. D-F12 = U87 cell culture medium. Slice = slice medium. NR = no significant response. ND = no data. Drugs used for slice experiments in bold. CUTOFF: at least ∼50 inhibition for CellTiter-Glo, minimum increase of CellTox Green signal for U87 of 200% (∼500% max). *for slice vs. cell medium (screen III only): red = increase in EC50 of more than 4 fold, blue = decrease in EC50 of more than 4 fold.

**Supplemental Table S2.** Secondary screen results with GBM8. 3 screen runs (I, II, and III) were performed with GBM8 cells. GSC = GBM8 glioma stem cell culture medium. Slice = slice medium. NR = no significant response. ND = no data. Drugs used for slice experiments in bold. CUTOFF: at least ∼50 inhibition for CellTiter-Glo, minimum increase of CellTox Green signal for GBM8 of 150% (100-200% max). *for slice vs. cell medium (screen III only): red = increase in EC50 of more than 4 fold, blue = decrease in EC50 of more than 4 fold.

| GBM8 EC50 [M] |  | Screen # | I | II | III | III | II | III | III |
| --- | --- | --- | --- | --- | --- | --- | --- | --- | --- |
|  |  | Assay | CellTiter-Glo | CellTiter-Glo | CellTiter-Glo | CellTiter-Glo | CellTox | CellTox | CellTox |
|  |  | Medium | GSC | GSC | GSC | Slice* | GSC | GSC | Slice* |
| Drug | Target | II,III max conc. |  |  |  |  |  |  |  |
| 17-AAG (Tanespimycin) | HSP (e.g. HSP90) | 10 uM | 29.0E-9 | 4.2E-9 | 4.2E-9 | 5.6E-9 | 4.8E-9 | 9.2E-9 | 4.7E-9 |
| Abiraterone (CB-7598) | P450 (e.g. CYP17) | 10 uM | NR | NR | NR | NR | NR | NR | NR |
| Afatinib | EGFR | 100 uM | 2.3E-6 | 534.8E-9 | 603.6E-9 | **10.0E-6** | 467.5E-9 | 463.9E-9 | **3.5E-6** |
| AMG 903 | Aurora Kinase | 10 uM | 441.9E-12 | 1.7E-9 | 3.6E-9 | **510.3E-9** | 1.7E-9 | 1.8E-9 | **50.0E-9** |
| APO866 (FK866) | O/metabolism | 10 uM | 44.6E-12 | 781.7E-12 | 466.4E-12 | **4.5E-9** | NR | NR | NR |
| BI 2539 | PLK | 10 uM | 3.0E-9 | 2.4E-9 | 4.1E-9 | 9.9E-9 | 2.1E-9 | 3.2E-9 | **72.5E-9** |
| Bortezomib (Velcade) | Proteasome | 10 uM | 1.6E-12 | 1.4E-9 | 3.2E-9 | **116.8E-9** | 1.43E-09 | 2.7E-9 | **59.2E-9** |
| Cisplatin | DNA/RNA Synthesis | 10 uM | ND | 66.3E-9 | 6.0E-6 | NR | 41.0E-9 | 360.8E-9 | NR |
| Danuserib (PHA-739358) | Aurora Kinase | 10 uM | 12.7E-9 | 7.6E-9 | 15.1E-9 | 33.7E-9 | 7.6E-9 | 14.4E-9 | 48.0E-9 |
| Dovitinib (TKI-258) | c-Kit | 10 uM | 121.8E-9 | 100.1E-9 | 137.5E-9 | 296.2E-9 | 165.2E-9 | 384.8E-9 | 330.4E-9 |
| Fingolimod (FTY720) | Bcr-Abl | 10 uM | 2.8E-6 | poor | 1.8E-6 | **52.2E-3** | 2.7E-6 | 1.2E-6 | **8.7E-3** |
| Geldanamycin | Autophagy/HSP90 | 10 uM | 597.1E-12 | 12.7E-9 | 5.5E-9 | 5.3E-9 | 66.1E-9 | 8.6E-9 | 5.7E-9 |
| GSK2126461 | mTOR | 10 uM | 16.9E-9 | 4.4E-9 | 7.6E-9 | 4.0E-9 | 66.4E-9 | 67.9E-9 | **8.4E-9** |
| Idarubicin HCl | Topoisomerase | 10 uM | 503.8E-12 | 29.1E-9 | 26.0E-9 | **218.1E-9** | NR | 24.0E-9 | **2.5E-6** |
| Imatinib | PDGFR/RTKi | 10 uM | 205.4E-9 | 391.2E-9 | 315.1E-9 | **14.5E-6** | NR | NR | **826.3E-9** |
| INK 128 (MLN0128) | mTOR | 10 uM | 5.2E-9 | 35.4E-9 | 96.0E-9 | 193.8E-9 | 1.6E-6 | 7.1E-6 | **1.2E-6** |
| Irinotecan | Topoisomerase | 10 uM | 566.2E-9 | 178.7E-9 | 210.1E-9 | 211.9E-9 | 142.8E-9 | 165.7E-9 | 190.0E-9 |
| Lomustine (CeeNU, CCNU) | O/Alkylator | 10 uM | 3.1E-6 | 95.6E-9 | 149.8E-9 | **793.4E-9** | 36.6E-9 | 90.5E-9 | **361.4E-9** |
| MLN2238 | Proteasome | 10 uM | 9.5E-9 | 17.3E-9 | 40.1E-9 | **4.3E-6** | 22.0E-9 | 41.4E-9 | **23.9E-3** |
| Mocetinostat (MGCD0103) | HDAC | 10 uM | 229.5E-9 | 122.1E-9 | 159.5E-9 | **1.2E-6** | 118.8E-9 | 131.0E-9 | **1.0E-6** |
| Motesanib | multiTK | 100 uM | ND | 27.7E-9 | 147.4E-9 | **30.7E-6** | ~1E-6 | 20.6E-6 | **677.0E-9** |
| Parthenolide | NFkB | 100 uM | ND | 1.7E-6 | 1.2E-6 | 2.1E-6 | 1.3E-6 | 1.2E-6 | 2.5E-6 |
| Pelitinib (EKB-569) | EGFR | 10 uM | 1.0E-6 | 701.3E-9 | 971.5E-9 | 2.5E-6 | 371.5E-9 | 394.8E-9 | 1.5E-6 |
| PI-103 | Autophagy/PI3K | 10 uM | 343.4E-9 | 727.3E-9 | 756.5E-9 | 1.1E-6 | 3.3E-6 | 3.3E-6 | 1.8E-6 |
| Ponatinib (AP24534) | Bcr-Abl | 30 uM | 35.0E-9 | 16.2E-9 | 15.4E-9 | **183.4E-9** | 141.8E-9 | 286.2E-9 | 231.9E-9 |
| Raltitrexed (Tomudex) | DNA/RNA Synthesis | 10 uM | 7.7E-9 | 26.6E-9 | 13.9E-9 | **234.1E-9** | 19.0E-9 | 11.5E-9 | 33.7E-9 |
| sorafenib | PDGFR/Tki | 30 uM | 39.8E-9 | 91.6E-9 | 82.6E-9 | **8.1E-6** | 3.5E-6 | 4.0E-6 | **18.6E-3** |
| temozolamide | Autophagy/Alkylator | 100 uM | 4.3E-6 | 271.6E-9 | 454.9E-9 | NR | 171.4E-9 | 442.4E-9 | 889.0E-9 |
| TW-37 | Bcl-2 | 10 uM | 3.9E-6 | 4.9E-6 | 3.2E-6 | 5.5E-6 | 518.3E-9 | NR | **21.2E-6** |
| vincristine | Autophagy/ Alkylator | 10 uM | 209.6E-12 | <300E-12 | <300E-12 | **446.4E-12** | <1E-9 | <300E-12 | **18.8E-9** |
| vorinostat | Autophagy/HDAC | 10 uM | 427.2E-9 | 373.6E-9 | 411.5E-9 | **1.9E-6** | 266.2E-9 | 370.3E-9 | **1.7E-6** |
| YM155 | E3 Ligase/survivin | 10 uM | 2.4E-9 | 5.5E-9 | 11.7E-9 | 5.1E-9 | 4.9E-9 | 19.5E-9 | 13.7E-9 |

